# High-fat feeding increases graft-derived glucagon output and impairs beta-cell secretion in transplanted human islets

**DOI:** 10.64898/2026.11.24.769013

**Authors:** Chunhua Dai, Katie C. Coate, Rebekah Brantley, Emily Acker, Amber Bradley, Abhilansh Pandey, Nandita Dey, Teri Doss, Radhika Aramandla, Adel R Eskaros, Ajay Singh, Mikaela Gay, Regina Jenkins, Jordan J. Wright, Leonard D. Shultz, Alvin C. Powers

## Abstract

Dysregulated glucagon secretion is central to the bihormonal disturbance that drives hyperglycemia in T2D, yet how a chronic obesogenic environment alters human alpha-cell function over time has not been directly examined. Models to specifically assess in vivo human alpha-cell function have not been possible because mouse and human glucagon are sequence-identical. We transplanted human islets (+I) into glucagon-knockout immunocompromised (GKO-NSG) mice, in which all circulating glucagon is graft-derived, and fed recipients a regular diet (RD+I) or a 60% high-fat diet (HFD+I) for 12 weeks. HFD+I recipients became obese, hyperlipidemic, hyperglycemic, and glucose intolerant, with elevated host mouse insulin indicating insulin resistance. Graft-derived human glucagon after glucose plus arginine was higher in HFD+I recipients at 4, 8, and 12 weeks. Fasting glucagon rose progressively between 4 and 12 weeks despite concurrent hyperglycemia and fasting human hyperinsulinemia, and graft endocrine composition and gross graft size were unchanged. Acute glucose-induced glucagon suppression persisted. Human beta-cells of the same grafts showed secretory dysfunction, so the impact on the two islet cell types diverged on a shared timeline. Conventional and multiplexed immunoimaging detected no accompanying changes in alpha-cell-enriched transcription factors and no change in human-derived immune, vascular, or matrix markers. Islet graft amyloid increased. In a contrasting insult, diphtheria toxin (DT) ablation of host beta-cells in GKO-NSG-DTR recipients produced severe hyperglycemia and reduced both hormones, suggesting that the direction of the human alpha-cell response may depend on the severity and nature of the insult. Inappropriate glucagon secretion is thus an early feature of the human islet response to diet-induced insulin resistance.

## 1 Introduction

Dysregulated glucagon secretion is a hallmark of type 1 diabetes (T1D) and T2D and is detectable in obese, insulin-resistant individuals before T2D is diagnosed.^1–6^ In diabetes, glucagon is inadequately suppressed after a meal, and fasting glucagon is inappropriately high, and both defects raise blood glucose by stimulating hepatic glucose production.^4,5,7–11^ Insulin deficiency magnifies that effect, which is the basis of the bihormonal view of diabetes.^6,7,10,12,13^ Too little glucagon is also harmful: the counter-regulatory response to hypoglycemia is markedly blunted in diabetes, most evidently in T1D.^14–16^ Alpha-cell dysfunction is therefore bidirectional, and it appears early: it contributes to impaired fasting glucose independently of insulin secretion, fasting glucagon predicts subsequent beta-cell decline, and abnormal glucagon secretion accompanies deteriorating glucose tolerance.^17–24^ How the human alpha cell responds to insulin resistance remains unclear. Because glucagon receptor agonists and antagonists now feature prominently in obesity and diabetes pharmacotherapy, how the human alpha-cell is regulated has become a therapeutic question as well as a physiologic one.^25–28^

Most of what we know about alpha-cell biology comes from rodents. However, rodent and human islets differ fundamentally in innervation^29,30^, vascularization^29,31,32^, cellular composition and arrangement^31,33^, hormone secretion^31,33^, mitogenic stimuli^34^, stress signaling^34,35^, lipid droplet accumulation^34^, and the distribution of islet-enriched transcription factors (TFs) such as MAFB.^32,36–40^ Alpha-cells themselves illustrate this point. They comprise less than 20% of mouse islet endocrine cells and are arrayed around the islet mantle, whereas they comprise more than 35% of human islet endocrine cells and are intermingled with non-alpha-cells throughout the islet, which implies different paracrine interactions.^31,33,41^ Rodent islets, therefore, have limitations for modeling human alpha-cell physiology or its dysfunction in T2D.

Studies of dispersed human alpha-cells have begun to define that failure directly. Relative to alpha-cells from donors without diabetes, glucagon exocytosis from T2D alpha-cells is blunted at low glucose, which is normally stimulatory, and paradoxically elevated at high glucose, which is normally inhibitory.^42–44^ These defects accompany reduced granule density, dysregulated ion channel activity, expression of genes characteristic of immature alpha-cells, and loss of paracrine inhibition by insulin and somatostatin.^42–44^ The in vitro phenotype is therefore net hyposecretion rather than the hypersecretion seen clinically, and glucagon secretion did not differ between isolated islets from donors with early-stage T2D and those without diabetes.^45^ Dysregulated glucagon secretion is therefore likely both alpha-cell-intrinsic and alpha-cell-extrinsic. Studying it in a system that preserves systemic inputs is critical.

Transplanted human alpha-cells have never been systematically studied in vivo in a chronic obesogenic environment. Ad libitum high-fat diet (HFD) feeding is a widely accepted model of obesity-induced insulin resistance and prediabetes.^46^ In rodents, 12-14 weeks of HFD elevates glucagon secretion from isolated islets and the perfused pancreas, and alters alpha-cell calcium signaling.^42,47^ Whether human alpha-cells respond in the same way is unknown. We previously transplanted human islets into NSG mice and found that a 12-week 60% HFD impaired nutrient-stimulated human insulin secretion and reduced graft MAFB and NKX6.1, whereas the same diet augmented insulin secretion and proliferation in host mouse beta-cells.^34^ Furthermore, human, but not mouse, islets accumulated superoxide, amyloid, and intracellular lipid.^34^ Since mouse and human glucagon are sequence-identical, human glucagon could not be measured in that model. GKO-NSG recipients lack endogenous glucagon, so all circulating glucagon is graft-derived and can be measured alongside human insulin in the same animal.^34,48^

Here, we show that a chronic obesogenic environment drives transplanted human alpha- and beta-cells in opposite secretory directions. In GKO-NSG recipients fed a 60% HFD for 12 weeks, graft-derived human glucagon was higher after combined glucose and arginine at 4, 8, and 12 weeks, and fasting glucagon rose progressively despite concurrent hyperglycemia and hyperinsulinemia. Acute glucose-induced suppression of glucagon nonetheless remained intact, so the dysregulation was secretagogue-selective rather than global. Beta-cells from the same grafts reproduced the secretory dysfunction previously reported in NSG recipients^34^, and the two cell types diverged on a shared timeline. To our knowledge, this is the first in vivo assessment of transplanted human alpha-cell function under long-term HFD and the first application of CODEX multiplexed imaging to human islet grafts. Inappropriate glucagon secretion emerges as an early feature of the human islet response to diet-induced metabolic stress, and GKO-NSG recipients enable concurrent in vivo assessment of human alpha- and beta-cell function.

## 2 Materials and Methods

### 2.1 Human islets

De-identified adult human islets (n = 6 preparations; Supplementary Table 1) from organ donors without diabetes were obtained from isolation centers within the Integrated Islet Distribution Program (IIDP; https://iidp.coh.org/), the Human Pancreas Analysis Program (HPAP)^49–52^ (https://hpap.pmacs.upenn.edu/), or the International Institute for the Advancement of Medicine (IIAM). Because de-identified tissue from deceased donors was used, these studies were not classified as human-subjects research. Islets were hand-picked to >95% purity and cultured in CMRL 1066 medium (5.5 mM glucose, 10% fetal bovine serum, 1% penicillin/streptomycin, 2 mM L-glutamine) at 37°C in 5% CO2 prior to transplantation. Donor characteristics are reported in Supplementary Table 1 in accordance with human-islet reporting guidelines.^53,54^ The six preparations used for the high-fat diet study were all from female donors (age 30-66 years; BMI 27.4-43.7 kg/m2), and four additional preparations (two from female and two from male donors) were used for the GKO-NSG-DTR studies. Islets from each preparation were divided between the two experimental groups within an experiment, so that every preparation contributed to both the RD+I and HFD+I groups, or to both the PBS and DT groups, and no preparation was allocated to a single group alone.

### 2.2 Mouse models

Immunocompromised glucagon-knockout NOD.Cg-Gcg ^em1^Dvs *Prkdc* ^scid^ *Il2rg ^tm1Wjl^* /DvsJ (GKO-NSG) mice were generated and characterized as previously described.^48^ Because these mice lack endogenous mouse glucagon, circulating glucagon in recipients of human islet grafts reflects human glucagon secreted by the graft, thereby enabling measurement of human alpha-cell function in vivo without confounding by host glucagon. Host beta-cell ablation was achieved using an NSG strain, NOD.Cg-Prkdcscid Il2rgtm1Wjl Tg(Ins2-HBEGF)6832Ugfm/Sz (NSG-RIP-DTR), which carries a rat insulin promoter-driven human HBEGF (diphtheria-toxin receptor, DTR) transgene and renders mouse beta-cells sensitive to diphtheria toxin (DT). DT sensitivity in this strain is conferred by the transgene.^34^ Originally generated in B6;CBA mice (provided by P.L. Herrera), the transgene was backcrossed onto the NSG background as previously described^34^ and crossed with GKO-NSG mice to generate NOD.Cg-Gcgem1Dvs Prkdcscid Il2rgtm1Wjl Tg(Ins2-HBEGF)6832Ugfm/Sz mice, referred to throughout as GKO-NSG-DTR. All animal procedures were approved by the Vanderbilt University Institutional Animal Care and Use Committee and were performed in accordance with institutional and federal guidelines. Mice were maintained by the Vanderbilt Division of Animal Care in group housing within a specific pathogen-free barrier facility on a 12 h light:12 h dark cycle with free access to water and food. For clarity, glucagon-knockout NSG recipients are referred to throughout as GKO-NSG and glucagon-knockout NSG recipients carrying the RIP-DTR transgene as GKO-NSG-DTR; recipients of a human islet graft are additionally denoted by +I.

### 2.3 Human islet transplantation and dietary intervention

Human islets (1,000 human islet equivalents [IEQ] per mouse) were transplanted under the kidney capsule of GKO-NSG or GKO-NSG-DTR recipients (12-20 weeks of age) using established procedures.^34,48^ Mice were anesthetized with isoflurane delivered by a SomnoFlo low-flow electronic vaporizer (Kent Scientific, Torrington, CT, USA). Anesthesia was induced with 3% isoflurane in an induction chamber for 3 min and maintained with 2% isoflurane via nose cone for the duration of surgery (approximately 30 min). For the HFD study, male GKO-NSG recipients were used (n = 23 RD+I and n = 23 HFD+I, derived from six donor preparations, with 3-5 mice per group per preparation); both male and female mice were studied in the GKO-NSG-DTR ablation experiments as indicated. The number of mice and the number of donor preparations contributing to each experiment are reported in the figure legends and in Supplementary Table 1. After a 2-week engraftment period, GKO-NSG recipients were assigned to regular diet (RD+I; Lab Diet 5001; 13.5% of calories from fat, 58% from carbohydrate, 28.5% from protein) or high-fat diet (HFD+I; Research Diets D12492; 60% of calories from fat, 20% from carbohydrate, 20% from protein) for 12 weeks. Non-fasting blood glucose and body weight were measured weekly. Blood glucose during serial monitoring was measured with a handheld glucometer (Contour next, Ascensia Diabetes Care US Inc), which reports values above 600 mg/dL as out of range. Values above 600 mg/dL are reported as 600 mg/dL and represent a lower bound. Plasma glucose at terminal assessments was measured by the Hexokinase/G6PDH method (Sigma, Cat. No 10737275001) which is not subject to this ceiling. Serum triglycerides (Thermo Scientific) and cholesterol (Thermo Scientific) were measured at 4, 8, and 12 weeks on diet. Intraperitoneal glucose tolerance tests (2 g/kg glucose) were performed after approximately 3, 7, and 11 weeks on diet, and glucose plus arginine stimulation tests (2 g/kg each) were performed after 4, 8, and 12 weeks on diet.

### 2.4 In vivo hormone secretion tests and amino acid measurements

Transplanted human alpha- and beta-cell function was assessed in vivo after a 5-6 h fast. For stimulation studies, mice were injected intraperitoneally with glucose alone or glucose plus arginine, and blood was collected before (0 min) and after (15 min) injection, as previously described.^34^ Glucagon was measured with the Mercodia Glucagon ELISA (cat. no. 10-1281-01), which the manufacturer reports to have little or no cross-reactivity with oxyntomodulin, mini-glucagon, GLP-1, GLP-2, or glicentin-related pancreatic polypeptide. Human insulin was measured with a human-specific insulin ELISA (Mercodia, cat. no. 10-1113-01) that does not cross-react with mouse insulin, and host mouse insulin was measured in a subset of animals after 4 weeks on diet with a mouse-specific insulin ELISA (Mercodia, cat. No 10-1247-01). Stimulated hormone values were obtained from a single 15 min sample. Total circulating amino acids (Sigma) were measured as an in vivo readout of hepatic glucagon action within the liver-alpha-cell axis.^48,55,56^

### 2.5 Graft histology, amyloid detection, and quantification

At study completion, mice were anesthetized with an intraperitoneal injection of ketamine (90-120 mg/kg) and xylazine (5-10 mg/kg) diluted in 0.9% sodium chloride. Graft-bearing kidneys were collected, after which mice were euthanized by cervical dislocation. Grafts were lightly PFA-fixed and processed for immunofluorescence as previously described.^16,34^ Primary and secondary antibodies and their dilutions are listed in Supplementary Table 2. Amyloid, a histopathological feature of human diabetic islets^57,58^, was detected by Thioflavin S staining and quantified relative to insulin-positive area as previously described.^34^ Images were acquired at 10x or 20x magnification on an Olympus confocal microscope and analyzed using HALO (Indica Labs) or ImageJ.

### 2.6 CODEX multiplexed spatial imaging

Graft sections were also processed and analyzed by CODEX multiplexed immunofluorescence to evaluate islet transcription factors, immune markers, endothelial markers, and extracellular matrix components in situ, as previously described.^45,59^ CODEX antibody panels and their dilutions are listed in Supplementary Table 3. All markers that passed validation in graft tissue are reported. Several antibodies in the panel, including CD45 and CD31, are human-specific and therefore detect only human-derived cells within the graft. Images were analyzed using HALO software (Indica Labs).

### 2.7 GKO-NSG-DTR ablation studies

To examine the impact of severe hyperglycemia on graft hormone secretion, human islets were transplanted into GKO-NSG-DTR recipients. After engraftment, mice received a single intraperitoneal injection of diphtheria toxin (DT; Biologicals, Cat. No 15047A1) at 5 ng per mouse in 300 uL sterile PBS, or an equal volume of PBS alone, and were monitored for 4 weeks (n = 12 PBS and n = 12 DT, derived from four donor preparations). DT was prepared and handled under institutionally approved biosafety procedures. Blood glucose was followed longitudinally, and fasting and stimulated plasma glucose, glucagon, and human insulin were measured. PBS-treated control mice received glucose plus arginine (2 g/kg each), as in the HFD study, because arginine alone does not raise blood glucose in this model. DT-treated mice were already severely hyperglycemic, so stimulated hormone secretion in that group was assessed after arginine alone (2 g/kg). Stimulated values in the two groups therefore reflect different secretagogue combinations, and this difference is considered in the interpretation. Total circulating amino acids and relative pancreas weight were measured as complementary readouts of impaired glucagon action and interrupted glucagon signaling.^55,56,60^ To assess whether human islet cells would be expected to be DT-sensitive in this model, HBEGF transcript levels were measured by quantitative RT-PCR in islets from nine human donors without diabetes and in islets from six NSG-RIP-DTR transgenic mice. RNA was isolated with (RNAqueous Micro Kit, Cat. No AM1931, Invitrogen), reverse transcribed with (High Capacity cDNA Reverse Transcription Kit, Cat. No 4368814, Appliedbiosystems), and amplified with (Taqman Universal PCR Master Mix, Cat. No 4304437, Appliedbiosystems); expression was normalized to *ACTB* and is expressed relative to the mean of the human preparations. Graft endocrine composition in PBS- and DT-treated recipients was quantified from whole-graft sections labeled for insulin, proglucagon, and somatostatin and is expressed as the percentage of hormone-positive cells per graft. Residual host beta-cells were quantified by counting insulin-positive cells per islet in pancreas sections from DT-treated male GKO-NSG-DTR mice.

### 2.8 Statistical analysis

Data are presented as mean ± SEM with individual data points shown in the figures. Comparisons between two groups were performed with Student t test or a nonparametric alternative, as appropriate. Within-test comparisons (0 versus 15 min) and comparisons among more than two groups used one-way or two-way ANOVA, as appropriate, with correction for multiple comparisons. In the high-fat diet study the same animals were sampled at 4, 8, and 12 weeks on diet, so measurements made at different timepoints in the same mouse are not independent. To test directly whether the fasting phenotype changed with duration of diet, fasting human glucagon and fasting human insulin in HFD+I recipients were compared across 4, 8, and 12 weeks by ANOVA with correction for multiple comparisons (Figure 2J, K). To assess whether the phenotype was consistent across islet preparations rather than driven by one or two, the 8-week stimulation data were additionally analyzed at the level of the donor preparation, with each symbol representing the mean of all mice transplanted with islets from a single preparation (Supplementary Figure 2A-C). The number of mice and the number of donor preparations are reported separately for each experiment. Blood glucose values recorded at the upper reporting limit of the handheld glucometer (600 mg/dL) are reported as 600 mg/dL and represent a lower bound. Statistical significance was defined as P < 0.05. GraphPad Prism 10 was used for graphing and statistical analyses.

## 3 Results

### 3.1 Chronic high-fat diet produces obesity, hyperlipidemia, hyperglycemia, and glucose intolerance in GKO-NSG human islet graft recipients

GKO-NSG mice enable direct measurement of graft-derived human glucagon (Figure S1). We first confirmed that our GKO-NSG recipients recapitulated the reported phenotypes of this model.^48^ Transplantation of 1,000 human IEQ (GKO+I) restored circulating glucagon and normalized blood glucose, glucose tolerance, total serum amino acids, relative pancreas weight, and alpha- and delta-cell hyperplasia relative to non-transplanted GKO mice and to NSG control mice (Figure S1). Because endogenous mouse glucagon is absent in this host, circulating glucagon in GKO+I recipients reflects human glucagon secreted by the graft, providing a direct readout of transplanted human alpha-cell function in vivo.

We next characterized the host response to chronic HFD-induced insulin resistance, used here to model the T2D milieu. Two weeks after transplantation, GKO-NSG recipients were assigned to a regular diet (RD+I) or HFD+I for 12 weeks (Figure 1A). HFD+I mice gained more body weight than RD+I controls, with significant divergence early in the dietary intervention that persisted through week 12 (Figure 1C). HFD+I recipients also developed hyperlipidemia, reflected by significantly elevated triglycerides and cholesterol at 4, 8, and 12 weeks on diet (Figure 1D, E). Non-fasting blood glucose was significantly elevated in HFD+I mice after 1 week of HFD exposure and remained elevated throughout the study (Figure 1F). Intraperitoneal glucose tolerance testing showed that HFD+I mice were glucose intolerant by 3 weeks and remained glucose intolerant at 7 and 11 weeks (Figure 1H-J). Host mouse insulin, measured after 4 weeks on diet, was higher in HFD+I recipients than in RD+I controls both at fasting and 15 min after glucose plus arginine (Figure 1G), confirming that recipients were hyperinsulinemic in the face of hyperglycemia, as expected with insulin resistance. Total circulating amino acids did not differ between RD+I and HFD+I recipients and were comparable to those of non-transplanted Gcg+/+ mice in both groups (Figure 1B), indicating that the arginine challenge was delivered against a comparable amino acid background in the two diet groups. These data validate that the HFD intervention produced the expected obesogenic, insulin-resistant, hyperlipidemic, and dysglycemic host environment in GKO-NSG recipients bearing human islet grafts.

**Figure 1.**
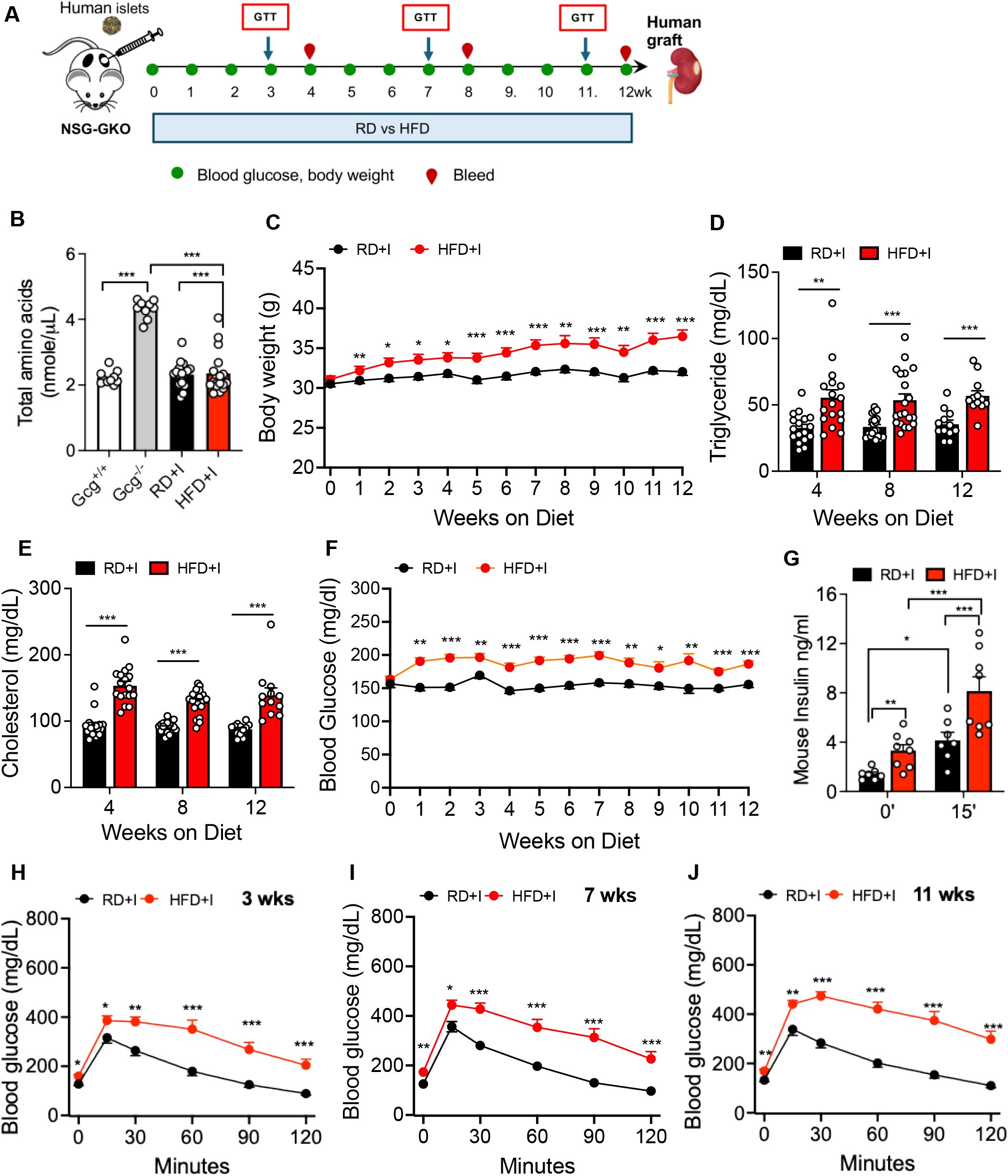
Chronic HFD produces obesity, hyperlipidemia, hyperglycemia, hyperinsulinemia, and glucose intolerance in GKO-NSG human islet graft recipients. (A) Schematic of the experimental timeline: human islets (I; 1,000 IEQ) were transplanted under the kidney capsule of GKO-NSG mice. After a 2-week engraftment period, mice were assigned to a regular diet (RD+I; 13.5% fat) or high-fat diet (HFD+I; 60% fat) for 12 weeks, after which grafts were collected for further analyses. In both designations, +I denotes a recipient of a human islet graft. Blood glucose and body weight (green dots) were measured weekly, blood (red dots) was collected at 4, 8, and 12 weeks, and ipGTTs were performed at 3, 7, and 11 weeks. (B) Total serum amino acids in non-transplanted Gcg+/+ and Gcg-/- (GKO) mice and in RD+I and HFD+I graft recipients. (C) Weekly body weights during the 12-week study in RD+I (black circles) and HFD+I (red circles) groups. (D) Serum triglycerides and (E) cholesterol at 4, 8, and 12 weeks on diet. Each dot represents a single mouse. RD+I, black bars; HFD+I, red bars. (F) Weekly non-fasting blood glucose during the 12-week study in RD+I (black circles) and HFD+I (red circles) groups. (G) Host mouse insulin at fasting (0 min) and 15 min after intraperitoneal glucose plus arginine after 4 weeks on diet (n = 7-8 mice per group from two donor preparations). (H-J) Intraperitoneal glucose tolerance tests after 3 (H), 7 (I), and 11 (J) weeks on diet in RD+I (black circles) and HFD+I (red circles) mice. Group sizes were 23 RD+I and 23 HFD+I mice derived from six human islet preparations (3-5 mice per group per preparation); the number of mice contributing to each panel is indicated by the individual data points where these are plotted. Data are mean ± SEM with individual points shown where applicable. *P < 0.05, **P < 0.01, ***P < 0.001.

### 3.2 Chronic HFD elicits a progressive increase in graft-derived human glucagon alongside human beta-cell secretory dysfunction

The human beta-cell response to chronic HFD in transplanted human islets has been characterized previously in NSG recipients^34^; the parallel human alpha-cell response, made measurable here by the GKO host, has not. To define that response and to compare it directly with the concurrent beta-cell response, we measured blood glucose, human glucagon, and human insulin at fasting and 15 min after combined glucose plus arginine stimulation at 4, 8, and 12 weeks of diet. At 4 weeks, we found that the blood glucose and glucagon responses to glucose plus arginine were significantly higher 15 min after stimulation in the HFD+I mice than in RD+I controls (Figure 2A-B). The concurrent human beta-cell measurements showed a distinct and opposite pattern: fasting human insulin was elevated in HFD+I mice, whereas RD+I mice increased human insulin after stimulation and HFD+I mice showed a reduction or failure of stimulated human insulin secretion (Figure 2C), confirming in the GKO-NSG background the fasting hyperinsulinemia with impaired stimulated secretion previously reported for HFD-exposed human beta-cell grafts.^34^ By 4 weeks, HFD was associated with increased graft-derived human glucagon after combined secretagogue stimulation, while human beta-cells showed the expected secretory failure.

**Figure 2.**
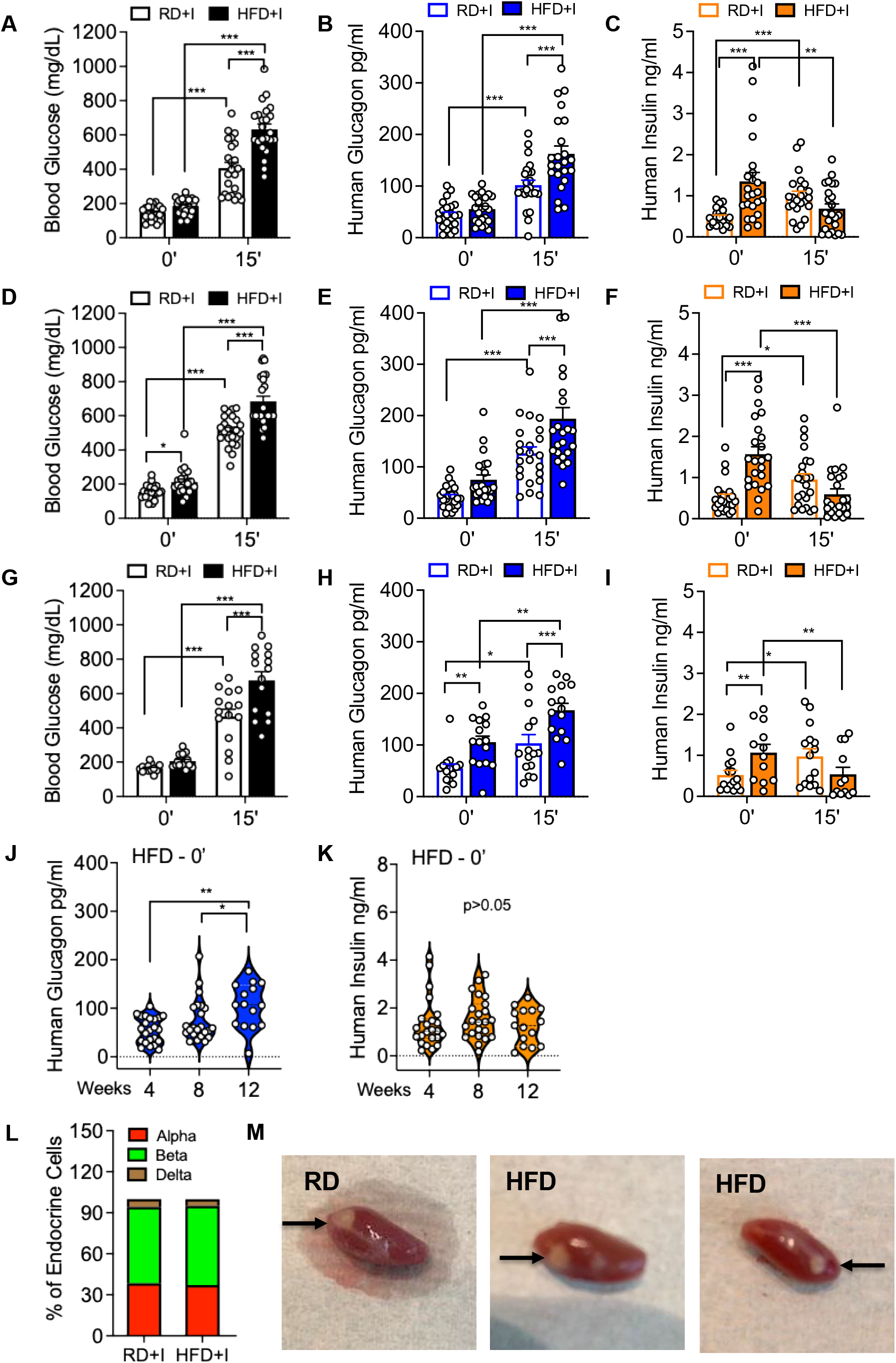
HFD produces divergent human alpha- and beta-cell secretory responses over time. (A, D, G) Blood glucose at fasting (0 min) and 15 min after intraperitoneal glucose plus arginine in RD+I (white bars) and HFD+I (black bars) mice at 4 (A), 8 (D), and 12 (G) weeks on diet. Each dot represents a single mouse. (B, E, H) Plasma human glucagon at fasting and 15 min after stimulation in RD+I (white bars) and HFD+I (blue bars) mice at 4 (B), 8 (E), and 12 (H) weeks. (C, F, I) Plasma human insulin at fasting and 15 min after stimulation in RD+I (white bars) and HFD+I (orange bars) mice at 4 (C), 8 (F), and 12 (I) weeks. (J, K) Fasting (0 min) plasma human glucagon (J) and human insulin (K) in HFD+I recipients at 4, 8, and 12 weeks on diet, shown as violin plots with individual mice and compared across time by repeated-measures analysis. (L) Endocrine cell composition of RD+I and HFD+I grafts. (M) Representative gross images of graft-bearing kidneys at retrieval after 12 weeks on diet, from an RD+I recipient and two HFD+I recipients transplanted with islets from the same donor preparation; arrows indicate the graft. The same animals were sampled at 4, 8, and 12 weeks. Data are from 23 RD+I and 23 HFD+I mice derived from six donor preparations; the number of mice contributing to each panel is indicated by the individual data points. Data are mean ± SEM with individual points shown. *P < 0.05, **P < 0.01, ***P < 0.001.

To assess whether the alpha-cell phenotype progressed, we measured secretion again at 8 and 12 weeks. At 8 weeks, we found that fasting blood glucose was significantly higher in the HFD+I group and remained higher after stimulation (Figure 2D), and human glucagon was again significantly higher after glucose plus arginine stimulation in HFD+I recipients (Figure 2E). Although fasting human glucagon did not reach significance at this time point, glucagon was not appropriately reduced despite concomitant fasting hyperglycemia and fasting hyperinsulinemia (Figure 2F), both of which should suppress alpha-cell activity further. By 12 weeks of HFD, while the elevated post-stimulation blood glucose response persisted (Figure 2G), we found that human glucagon was now significantly elevated in HFD+I mice both at fasting and after glucose plus arginine stimulation (Figure 2H), while the human insulin secretory defect persisted, with elevated fasting insulin and impaired stimulated insulin secretion (Figure 2I). Notably, endocrine cell composition of the graft was similar between diet groups (Figure 2L), and grafts retrieved from RD+I and HFD+I recipients transplanted with islets from the same donor preparation were of comparable gross size (Figure 2M), arguing against an increase in graft alpha-cell number as the explanation for the higher circulating glucagon. To evaluate the progression of the fasting phenotype, we compared fasting human glucagon in HFD+I recipients across 4, 8, and 12 weeks and found that it rose significantly with duration of diet, whereas fasting human insulin did not change over the same interval (Figure 2J, K). Because six donor preparations were distributed across a larger number of recipient mice, we also analyzed the 8-week data at the level of the donor preparation, with each symbol representing the mean of all mice transplanted with islets from a single preparation; the higher stimulated human glucagon and the higher fasting human insulin in HFD+I recipients were both reproduced in this analysis (Supplementary Figure 2A-C), indicating that the phenotype was consistent across preparations. These findings show that HFD exposure was associated with a progressive deterioration of fasting glucagon control and sustained augmentation of stimulated glucagon secretion by human alpha-cells.

### 3.3 Acute glucose-induced suppression of human glucagon secretion is preserved as fasting glucagon rises

To test whether the augmented glucagon phenotype reflected a complete loss of glucose-dependent suppression, we examined blood glucose and human glucagon at 0 and 15 min during intraperitoneal glucose tolerance testing across successive time points, with glucose as the sole secretagogue (Figure S2D-I). We found that at 3 weeks the acute glycemic excursion 15 min after glucose was significantly greater in HFD+I mice, yet human glucagon was appropriately suppressed 15 min after glucose in both groups (Figure S2D, E). By 7 and 11 weeks, fasting human glucagon was significantly higher in HFD+I recipients, but glucose still suppressed human glucagon at 15 min in HFD+I grafts (Figure S2F-I). These data indicate that HFD+I alpha-cells retained the capacity for acute glucose-induced glucagon suppression even as fasting glucagon rose. Because HFD+I mice began from a higher fasting glucagon and mounted a larger glycemic excursion, and therefore received a stronger suppressive signal, comparable absolute suppression in the two groups is likely to understate a relative impairment of the HFD+I alpha-cell, consistent with the inappropriately elevated fasting glucagon observed despite concurrent hyperglycemia and hyperinsulinemia described above. The phenotype of increased glucagon was most evident under combined glucose plus arginine stimulation and in the fasting state, closely paralleling the selective, secretagogue-dependent alpha-cell dysregulation described in human diabetes.

### 3.4 The human alpha-cell secretory phenotype occurs without loss of alpha-cell MAFB or misexpression of NKX6.1

To determine whether the divergent human alpha- and beta-cell secretory phenotypes were accompanied by changes in key transcription factors that regulate islet cell identity and hormone output, we quantified MAFB and NKX6.1 in graft alpha- and beta-cells after 12 weeks of diet. We selected these two candidate factors because they are the only islet-enriched transcription factors previously reported to be altered in this xenograft paradigm^34^, because MAFB maintains adult human alpha-cell identity and glucagon secretion^61^, and because NKX6.1, which is enriched in beta-cells, is an essential regulator of beta-cell identity and function.^62^ We found that the proportion of MAFB-positive beta-cells was significantly reduced in HFD+I grafts compared with RD+I controls (Figure 3A-B), and that NKX6.1 followed the same pattern (Figure 3E-F). This confirms the beta-cell transcription factor downregulation reported previously in NSG recipients.^34^ Conversely, we found that neither factor differed between diet groups in glucagon-positive alpha-cells: alpha-cell MAFB was not reduced, and NKX6.1 was not induced, in HFD+I grafts (Figure 3C-D, G-H). This indicates that the augmented human glucagon secretion arose without a detectable change in the abundance or nuclear localization of either factor in alpha-cells, and it directed us to look for other correlates of the alpha-cell phenotype.

**Figure 3.**
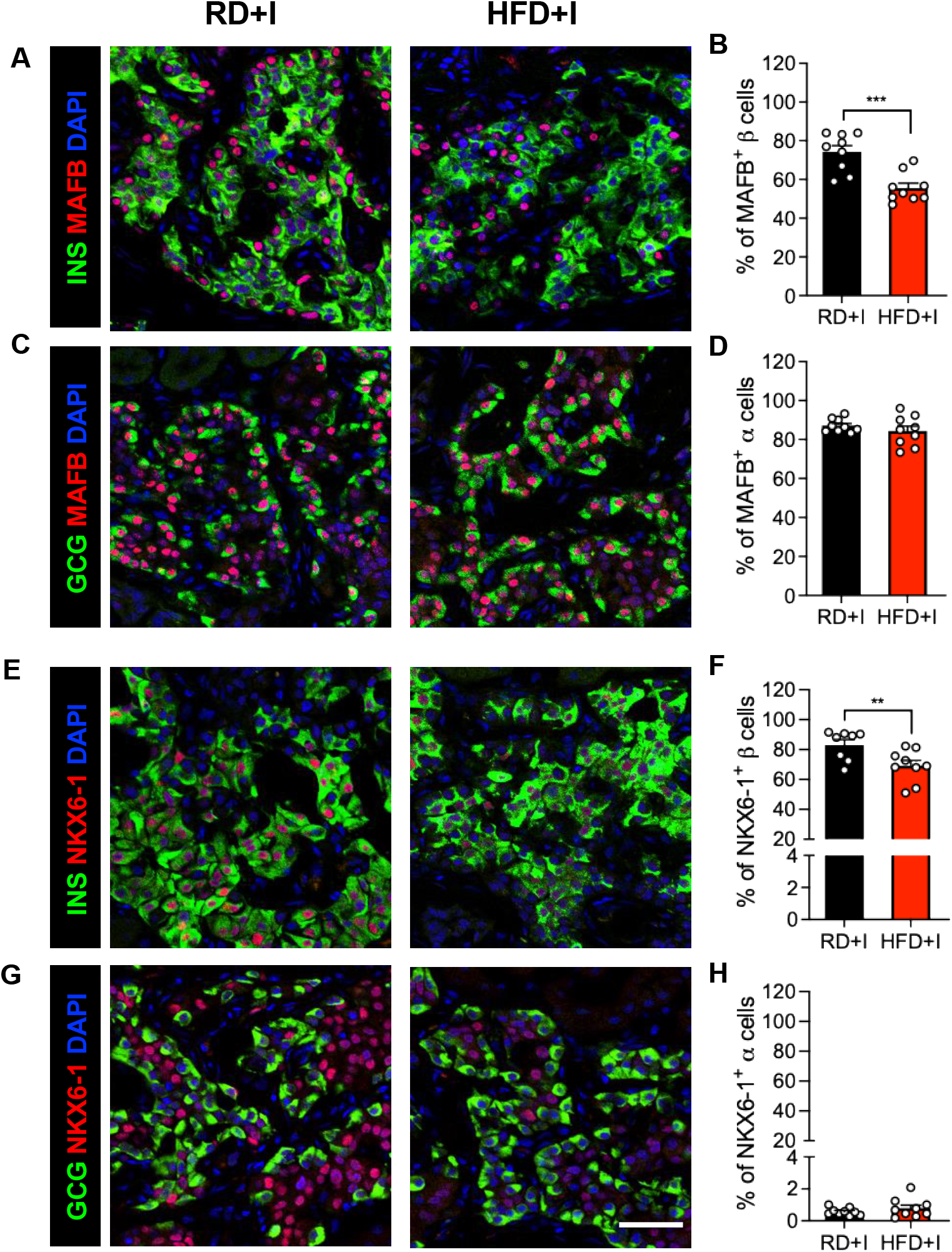
HFD reduces MAFB- and NKX6.1-positive beta-cells but not alpha-cells in human islet grafts. Representative immunofluorescent images of human grafts from RD+I (black bars) and HFD+I (red bars) mice. (A, B) INS (beta-cells, green), MAFB (transcription factor, red), and DAPI (nuclei, blue), with quantification of the percentage of MAFB-positive beta-cells. (C, D) GCG (alpha-cells, green), MAFB (red), and DAPI (blue), with quantification of the percentage of MAFB-positive alpha-cells. (E, F) INS (green), NKX6.1 (transcription factor, red), and DAPI (blue), with quantification of the percentage of NKX6.1-positive beta-cells. (G, H) GCG (green), NKX6.1 (red), and DAPI (blue), with quantification of the percentage of NKX6.1-positive alpha-cells. Scale bar = 40 µm for all images in this figure. Data are mean ± SEM with individual points shown. **P < 0.01, ***P < 0.001.

### 3.5 The human alpha- and beta-cell secretory phenotypes occur with increased graft amyloid but without changes in human-derived immune, vascular, or matrix markers

To determine whether the secretory changes were accompanied by the broader islet remodeling processes that are characteristic features of diabetic islets^45,63,64^, we performed CODEX multiplexed imaging of human islet grafts, selecting markers that report on processes previously implicated in alpha- and/or beta-cell dysfunction in diabetes. We found that, beyond the beta-cell MAFB and NKX6.1 reductions described above, no additional islet transcription factors differed significantly between diet groups: MAFA-positive beta-cells and ARX-positive alpha-cells were similar between groups (Figure S3A-D), and PAX6, PDX1, ISL1, IRX2, and NEUROD1 were unchanged in alpha- or beta-cells (Figure S3E-I). This indicates that, among the factors included in our panel, the transcription factor phenotype was restricted to reduced MAFB and NKX6.1 in beta-cells as we reported previously^34^, and is consistent with MAFB being among the islet-enriched transcription factors most sensitive to metabolic stress.^65,66^

We also found that the human-derived immune, vascular, and stromal markers in our panel were largely unchanged. The macrophage and antigen-presenting markers CD68, CD163, MMR, and HLA-DR, the macrophage marker IBA1, the pericyte marker MCAM, and the endothelial markers CD31 and SELP were not significantly altered by diet, and CD45-positive immune area relative to endocrine area was unchanged (Figure S4C-D; Figure S5A-G, M). Laminin, collagen IV, collagen VI, vimentin, and keratin were also unchanged, and collagen I trended higher in HFD+I grafts without reaching significance (Figure S5H-L, N; P = 0.0565 for collagen I). The one exception was islet amyloid, a characteristic feature of the T2D islet, which increased from an average of 3.8% of insulin-positive area in RD+I grafts to approximately 8.2% in HFD+I grafts (Figure S4A-B). This confirms in the GKO-NSG background the graft amyloid pathology previously reported in HFD-exposed human islets.^34^ Several antibodies in this panel, including CD45 and CD31, are human-specific and therefore report on the human-derived compartment of the graft rather than on host mouse immune or endothelial cells. We have previously shown that human islet grafts in this model retain a substantially human-derived vasculature after long-term engraftment^34^, so CD31 is informative about graft vessels; host myeloid cells and host endothelium, however, were not assessed here. Together, this indicates that, apart from amyloid deposition, the prominent human alpha- and beta-cell secretory phenotypes emerged without overt evidence of the human-derived islet inflammation, vascular rarefaction, or fibrotic matrix remodeling that characterize diabetic islets^45,63^, suggesting that the functional dysregulation of the graft precedes, or occurs independently of, these structural hallmarks over the timeframe studied.

### 3.6 Chronic extreme hyperglycemia in the GKO-NSG-DTR model is associated with reduced human alpha- and beta-cell hormone secretion

Having characterized human alpha-cell function under a chronic HFD, we next used the GKO-NSG-DTR model to assess how transplanted human alpha-cells (and beta-cells) respond to a more severe metabolic insult. In this model, DT-mediated ablation of host pancreatic beta-cells produces sustained hyperglycemia, allowing us to assess human graft hormone secretion under extreme hyperglycemia (Figure 4A). Because the DTR system depends on a human HBEGF transgene and human cells express HBEGF endogenously, we first asked whether human islet cells would themselves be expected to be DT-sensitive. HBEGF transcript levels in islets from nine human donors were approximately 20-fold lower than in islets from NSG-RIP-DTR transgenic mice (Figure 4B), consistent with DT sensitivity in this model being conferred by the transgene rather than by endogenous human HBEGF, and with our previous finding that DT ablates mouse but not human beta-cells in this transplantation setting.^34^ We found that DT treatment, but not PBS, led to a progressive increase in blood glucose, with values reaching the 600 mg/dL upper reporting limit of the handheld glucometer by approximately day 7 and remaining at or near that limit through the 4-week follow-up (Figure 4C; values recorded at the upper reporting limit of the handheld glucometer are reported as 600 mg/dL and represent a lower bound, whereas terminal plasma glucose was measured by a different method and is correspondingly higher). At the terminal assessment, DT-treated mice showed severe hyperglycemia together with reduced fasting human glucagon and insulin compared with PBS-treated controls (Figure 4D-F). This indicates that, under chronic extreme hyperglycemia, human alpha-cells no longer exhibit the relative fasting hyperglucagonemia seen under the milder HFD insult and instead show reduced hormone output alongside the graft beta-cell, a pattern we interpret as concurrent dysfunction of both alpha- and beta-cells in the setting of severe hyperglycemia, although these experiments do not establish the underlying mechanism.

**Figure 4.**
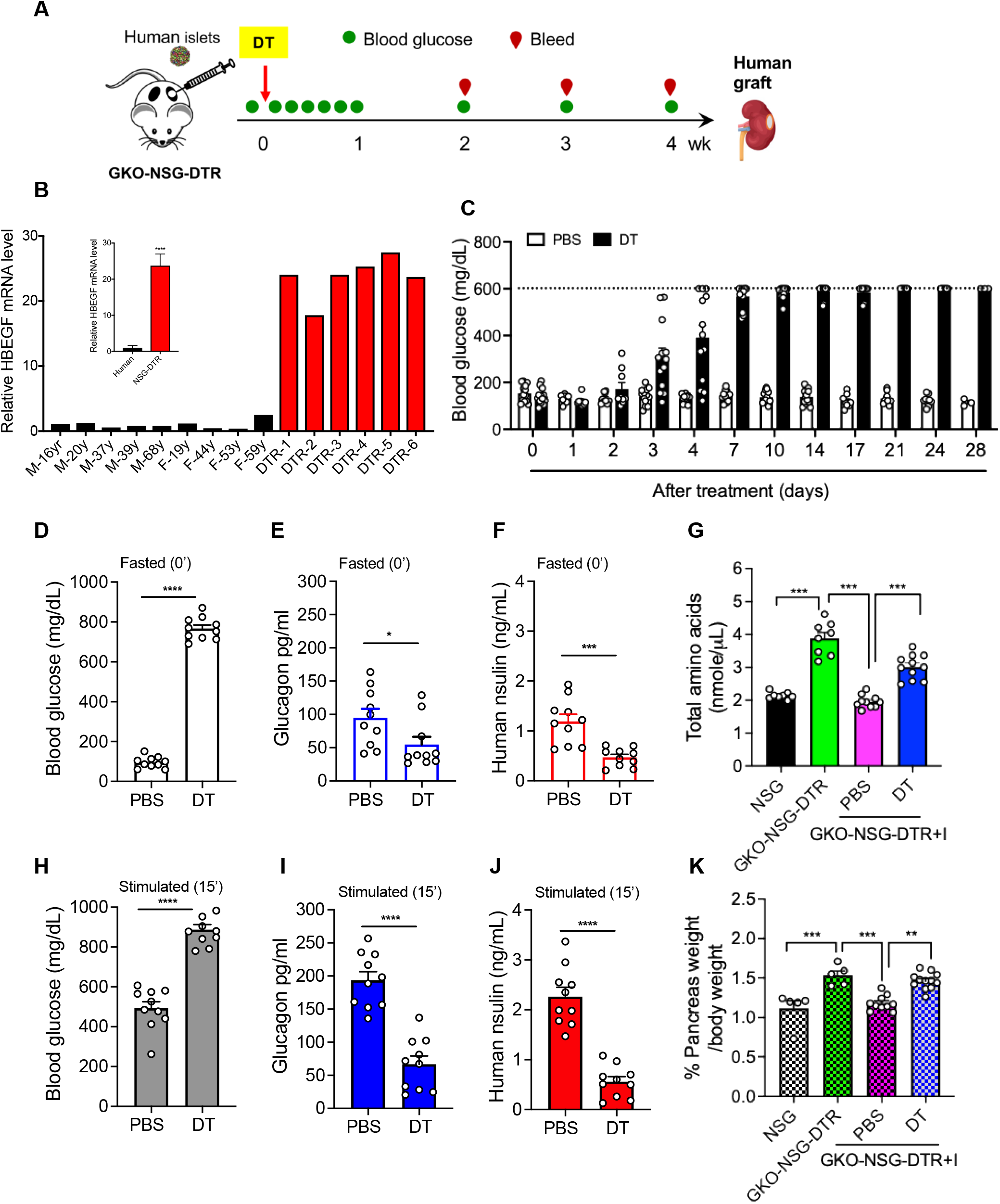
Chronic extreme hyperglycemia in the GKO-NSG-DTR model is associated with reduced human alpha- and beta-cell hormone secretion. (A) Schematic of the experimental design. Human islets were transplanted into GKO-NSG-DTR mice, followed by a single intraperitoneal injection of diphtheria toxin (DT; 5 ng per mouse) or vehicle (PBS), serial blood glucose monitoring (green dots), blood collection (red dots), and graft retrieval at the study endpoint. (B) HBEGF transcript levels measured by quantitative RT-PCR in islets from nine human donors without diabetes (black bars; donor sex and age indicated) and from six NSG-RIP-DTR transgenic mice (red bars), expressed relative to the mean of the human preparations. Inset, grouped comparison of human versus NSG-RIP-DTR islets. (C) Blood glucose over 28 days after PBS (white bars) or DT (black bars) treatment, measured with a handheld glucometer; the dotted line marks the 600 mg/dL upper reporting limit of the meter, and values at this limit are plotted at 600 mg/dL. (D-F) Fasting plasma glucose (D), human glucagon (E), and human insulin (F) in PBS- and DT-treated mice. (G) Total circulating amino acids in NSG, GKO-NSG-DTR, and human islet-transplanted GKO-NSG-DTR (GKO-NSG-DTR+I) mice treated with PBS or DT. (H-J) Stimulated plasma glucose (H), human glucagon (I), and human insulin (J) 15 min after glucose plus arginine in the PBS group or after arginine alone in the DT group. (K) Relative pancreas weight in the same groups shown in (G). n = 12 PBS and 12 DT mice derived from four donor preparations. Data are mean ± SEM with individual points shown. *P < 0.05, **P < 0.01, ***P < 0.001, ****P < 0.0001.

To probe stimulated secretion under this condition, we measured hormone output after arginine stimulation alone (because glucose levels were already markedly increased). We found that DT-treated mice remained more hyperglycemic and showed markedly lower arginine-stimulated glucagon and human insulin secretion compared to PBS controls (Figure 4H-J). Because PBS-treated controls received glucose plus arginine while DT-treated mice received arginine alone, the stimulated comparison is between different secretagogue combinations. DT treatment also increased total circulating amino acids and relative pancreas weight (Figure 4G, K), consistent with the loss of graft-derived glucagon action.^48,55,56,60^ This indicates that chronic extreme hyperglycemia is associated with failure of both human alpha- and beta-cell hormone secretion, in contrast to the increased graft-derived human glucagon elicited by 12 weeks of HFD, and suggests that the direction of the human alpha-cell response may depend on the severity and nature of the metabolic insult.

To determine whether the reduction in both graft hormones reflected loss of graft endocrine cells, we examined the grafts at retrieval. Graft-bearing kidneys from PBS- and DT-treated recipients were grossly similar (Figure S7A), and whole-graft sections labeled for insulin, proglucagon, and somatostatin showed comparable graft cross-sectional area with abundant endocrine cells in both groups (Figure S7B). Quantification of graft endocrine composition showed a modest reduction in the proportion of alpha-cells and a reciprocal increase in the proportion of beta-cells in DT-treated recipients, with delta-cells unchanged (Figure S7C). The graft therefore retained a substantial beta-cell complement in DT-treated animals even though human insulin secretion was markedly reduced, and graft alpha- and beta-cell proportions moved in opposite directions, which is not the pattern expected from non-selective DT-mediated killing of human graft cells. Graft endocrine mass could not be measured in this model, because a graft beneath the kidney capsule cannot be dissected free of host tissue and weighed in the way that pancreas weight is used to calculate islet cell mass. A contribution of altered graft cell number to the reduced hormone output therefore cannot be entirely excluded.

During validation of the model, the glycemic response to DT differed by sex. Male GKO-NSG-DTR mice became markedly hyperglycemic by day 3 and remained so throughout follow-up (Figure S6A), whereas female mice responded heterogeneously, including non-responders that maintained near-normal glucose and responders that developed variable, often delayed, hyperglycemia (Figure S6B-C). The basis for this difference is unknown and should be considered when designing future ablation studies in GKO mice. In both sexes, immunofluorescence for insulin, proglucagon, and somatostatin in host pancreas confirmed near-complete loss of host beta-cells after DT (Figure S6D, E): among 28 islets examined across 5 DT-treated male mice, 26 contained two or fewer insulin-positive cells and none contained more than five.

## 4 Discussion

Whether transplanted human alpha-cells secrete glucagon inappropriately in a sustained obesogenic host environment had not been directly assessed, because conventional xenograft hosts contribute their own glucagon to the circulation. The GKO-NSG model removes that constraint^48^, allowing graft-derived human glucagon and insulin to be measured in the same animals across 12 weeks of HFD, during which the host became obese, hyperlipidemic, dysglycemic, and hyperinsulinemic. Under these conditions the two cell types diverged. Human alpha-cells released excess glucagon after glucose plus arginine and developed progressive fasting hyperglucagonemia despite concurrent hyperglycemia and hyperinsulinemia, whereas human beta-cells lost regulated insulin secretion early and persistently, lost MAFB- and NKX6.1-positive cells, and accumulated amyloid, reproducing what we reported previously in NSG recipients.^34^ None of the alpha-cell transcription factors we examined changed over the same interval, although a targeted panel cannot exclude broader remodeling of alpha-cell identity programs. Long-term HFD therefore produced opposite secretory phenotypes in transplanted human alpha- and beta-cells: progressively augmented glucagon secretion and early, persistent beta-cell failure.

The pattern we observed, exaggerated glucagon release to glucose plus arginine alongside a failure to suppress fasting glucagon, mirrors a dissociation recognized in human diabetes for more than half a century. Studies by Unger and colleagues^10–12,67^ established that the diabetic alpha-cell retains intact, or even amplified, responses to arginine and other amino acids while losing glucose-dependent regulation. That dissociation anchored the bihormonal hypothesis, in which relative hyperglucagonemia during insulin resistance or insulin deficiency worsens hyperglycemia.^13^ Transplanted human alpha-cells reproduce this classic signature in response to a defined dietary insult, in a setting where the human origin of the glucagon is unambiguous.

The present work extends the NSG-HFD paradigm established by Dai et al.^34^ In that study, transplanted human islets challenged with chronic hyperglycemia, chronic insulin resistance, or acute insulin resistance responded differently from host mouse islets under the same conditions, and HFD-exposed grafts developed the beta-cell phenotype that we reproduce here on a glucagon-deficient background. The advance is the ability to measure human glucagon, which converts a beta-cell-focused stress paradigm into a bihormonal one.

Of the transcription factors in our panel, only MAFB and NKX6.1 changed, and only in beta-cells, where the proportion of positive cells fell by up to 25%. This partial downregulation is consistent with MAFB being among the earliest factors reduced when adult islet cells become dysfunctional.^34,66^ Because adult rodent beta-cells do not express MAFB^68^, such stress-sensitive downregulation can be detected only in human or non-human primate islets. MAFB knockdown in adult human islets modestly reduces insulin secretion^61,69^, so a partial reduction in MAFB, together with the concurrent reduction in NKX6.1, may contribute to the secretory defect observed here.

In alpha-cells, augmented glucagon release emerged by 4 weeks and persisted through 12 weeks, with no detectable changes in any transcription factors we examined (Figures 3 and S3). Because secretion increased rather than fell, the capacity to secrete glucagon was retained. We did not measure glucagon synthesis or intracellular glucagon content, so we cannot determine whether hormone production also changed, and unchanged transcription factor abundance does not exclude altered transcription factor activity or altered expression of target genes.

Nutrient sensing, electrical activity, paracrine input, and receptor responsiveness can each shift without a measurable change in transcription factor protein; we offer these as possible explanations for the phenotype rather than as established mechanisms, since none was directly tested here.^70^ Such dissociation between identity factor abundance and secretory competence is not peculiar to our model: single-cell analysis of T2D human alpha-cells found the cells expressing the highest levels of MAFB, ARX, and related factors to be the most functionally impaired.^42^ Reduced alpha-cell MAFB is itself reported in longer-standing T1D and T2D^16,66,71^, or after pharmacological ER stress in stem cell-derived alpha-cells^65^, so a longer or more severe insult than 12 weeks of HFD may be required before alpha-cell identity programs are measurably remodeled.

The DT ablation experiment models severe glucotoxicity and, by exposing human alpha-cells to an insult more extreme than HFD, delineates how sustained hyperglycemia affects human glucagon secretion. DT treatment produced severe hyperglycemia, reduced fasting and arginine-stimulated human glucagon and insulin, elevated circulating amino acids, and increased relative pancreas weight, the last two consistent with disruption of the liver alpha cell axis.^56,60^ The glucagon result is informative because it separates two possibilities. If hyperglycemia suppressed the alpha-cell mainly through paracrine insulin and somatostatin, the concurrent failure of graft insulin secretion should have released the alpha-cell and permitted, or even amplified, glucagon output. Human glucagon instead fell below that of PBS-treated controls, at fasting and under arginine stimulation. An alternative explanation, that DT acted directly on the human graft, is unlikely: HBEGF transcript levels in human islets are far below those in DTR transgenic islets (Figure 4B), graft endocrine content and gross graft size were preserved (Figure S7A-C), and graft alpha- and beta-cell proportions moved in opposite directions rather than falling together. We therefore favor dysfunction of both human alpha- and beta-cells at this degree of hyperglycemia. We cannot exclude a cell-intrinsic inhibitory response to severe hyperglycemia that is preserved rather than pathological, but either reading places the two models at different points on a stress continuum, with HFD associated with inappropriately increased graft-derived glucagon and extreme hyperglycemia with reduced secretion of both hormones.

In vivo, elevated fasting glucagon in individuals without diabetes predicts subsequent decline in beta-cell function, and impaired glucose suppression of glucagon contributes to impaired fasting glucose independently of any defect in insulin secretion.^17,21^ Over three years, that suppression deteriorates specifically in those progressing to impaired glucose tolerance or T2D, alongside declining beta-cell function.^22^ Meta-analysis confirms inadequate glucagon suppression during oral glucose tolerance testing in prediabetes, and in established T2D, impaired suppression tracks insulin resistance rather than insulin secretory failure.^19,20^ Alpha-cell dysfunction is therefore detectable before diabetes is diagnosable and is not simply a consequence of beta-cell failure. Our HFD data follow that trajectory, with glucagon becoming inappropriate for the prevailing glucose and insulin concentrations.

Ex vivo, the evidence is less consistent and, in some respects, points in the opposite direction. Isolated islets from donors with T2D show impaired glucagon granule exocytosis together with loss of paracrine restraint by insulin and somatostatin^44^, and a heterogeneous exocytotic defect linked to alpha-cell maturation state^42^, whereas islets from donors with short-duration T2D (mean 3.5 years) secreted glucagon indistinguishably from non-diabetic controls despite substantially impaired insulin secretion.^45^

Three explanations are compatible with these observations. First, alpha-cell dysfunction may require longer disease duration than the mean 3.5 years^45^, paralleling the delayed changes in alpha-cell identity discussed above. Second, standard ex vivo media may fail to unmask an existing defect if it lacks the physiologic complement of amino acids and other secretagogues needed to expose it. Third, the isolated islet may lack the systemic inputs that shape alpha-cell secretion in vivo, including hepatic glucagon resistance and the ensuing hyperaminoacidemia, incretin signals, sympathetic tone, and circulating metabolites. Transplantation data are most consistent with the third: T2D islets in GKO-NSG hosts secrete excess glucagon under basal and hypoglycemic condition^48^, and here non-diabetic islets in the same host background did so once the host became insulin-resistant, with dysregulation driven by donor disease in one case and by host diet in the other. What remains unresolved is how much of that dysfunction is alpha-cell intrinsic, how much is paracrine, and how much is imposed by the systemic environment. Separating them will require amino acid-inclusive secretion protocols applied to early-T2D islets both ex vivo and after transplantation into a glucagon-deficient host, an experiment not yet done.

A candidate mechanism, not tested here, is loss of paracrine restraint on the alpha-cell. Rodent HFD models show that reduced somatostatin signaling and alpha-cell somatostatin resistance blunt glucose-induced suppression of alpha-cell activity^42,47^, human islets depend on paracrine interactions to set the glycemic set point^72^, and the human T2D alpha-cell loses inhibition by insulin and somatostatin as somatostatin receptor availability falls.^44^ Our HFD+I grafts released glucagon despite elevated glucose and fasting human insulin, so the signals that normally restrain the alpha-cell were ineffective. Because the beta-cell defect appeared by 4 weeks and preceded fasting hyperglucagonemia at 12 weeks, we hypothesize that paracrine insulin restraint is lost progressively.

Multiplexed spatial proteomics, principally imaging mass cytometry, has previously mapped the native human pancreas in T1D^73^ and T2D.^45,74^ Applying CODEX to endpoint islet graft material opens the way to larger panels covering paracrine receptors, stress responses, vascular remodeling, and cell-cell neighborhoods. That the graft immune, vascular, and matrix compartments remained largely intact while secretory behavior changed markedly also fits the possibility raised above, that the systemic environment contributes substantially to the alpha-cell phenotype.

Several limitations qualify these conclusions: (i) We transplanted islets from donors without diabetes into immunodeficient mice, so the graft environment differs from the native human pancreas; (ii) The GKO background removes host glucagon, oxyntomodulin, and glicentin^48^ but leaves other host hormones and metabolic signals that may shape graft behavior; (iii) The study is descriptive. We report associations between diet, secretory phenotype, and a panel of candidate markers selected from prior work; however, we did not manipulate these markers, so no causal relationship can be inferred. Unbiased approaches such as single-cell transcriptomics or proteomics would be required to define graft-wide molecular changes and heterogeneity; (iv) We measured hormone secretion serially but analyzed transcription factors and CODEX markers only at the endpoint. Immunodetection reports protein abundance and localization rather than transcriptional activity or target gene output, so graded changes in transcription factor function, altered cofactor availability, or altered expression of target genes would not be captured. Furthermore, several of the antibodies used for CODEX, including CD45 and CD31, are human-specific, so they detect only the transplanted human cells and not host mouse immune or endothelial cells. Temporally matched graft analyses will be needed to identify early molecular correlates of alpha-cell dysregulation; (v) We did not measure somatostatin secretion, alpha-cell receptor expression, beta-cell glucagon receptor signaling, hepatic glucagon signaling, glucagon synthesis or intracellular glucagon content, or circulating GLP-1, any of which could contribute to the phenotype and to the sex-dependent variability seen in the DT model; (vi) Stimulated secretion was assessed from a single 15 min sample, so a shifted or prolonged secretory peak cannot be distinguished from a true increase in secretion; (vii) Graft endocrine mass could not be measured because the graft cannot be excised and weighed as the pancreas is when islet cell mass is calculated. While graft endocrine composition and gross graft size were similar between diet groups, a contribution of graft alpha-cell number to the increase in circulating graft-derived glucagon cannot be excluded, and for this reason we describe the phenotype as increased graft-derived glucagon rather than as hypersecretion per cell; (viii) The HFD study used male recipients only, and all six islet preparations used for it came from female donors, several with a BMI in the obese range, so islets from insulin-resistant donors may already have been adapted.

Within those constraints, this work joins an established human beta-cell xenograft stress paradigm to clinical evidence that alpha-cell dysfunction precedes and accompanies progression toward T2D. The GKO-NSG model and the CODEX graft-imaging workflow now make it possible to ask how paracrine signaling, hepatic glucagon resistance, and systemic metabolic signals converge on human alpha-cell programs during metabolic disease, and whether restoring appropriate glucagon suppression can slow that progression.

---

**Figure S1. Characterization of the GKO-NSG model and its normalization by a human islet graft.** Comparison of NSG (white), GKO (black), and GKO+I (gold) mice for (A) serum glucagon, (B) blood glucose, (C) intraperitoneal glucose tolerance, (D) total serum amino acids, and (E) relative pancreas weight. (F) Representative immunofluorescence images of pancreatic islets from NSG (left panel), GKO (middle panel), and GKO+I (right panel) mice labeled for insulin (INS, green), pro-glucagon (Pro-GCG, red), and somatostatin (SST, blue). Scale bar = 50 µm for all images in panel F. Data are mean ± SEM with individual points shown. ***P < 0.001, ****P < 0.0001.

**Figure S2. Donor-level analysis of the 8-week secretory phenotype, and blood glucose and human glucagon responses during intraperitoneal glucose tolerance testing across time.** (A-C) Blood glucose (A), plasma human glucagon (B), and plasma human insulin (C) at fasting (0 min) and 15 min after intraperitoneal glucose plus arginine after 8 weeks on diet, analyzed at the level of the islet preparation: each symbol is the mean of all mice transplanted with islets from one of six donor preparations. RD+I, open bars; HFD+I, filled bars. (D-I) Blood glucose (D, F, H) and plasma human glucagon (E, G, I) at fasting (0 min) and 15 min after intraperitoneal glucose alone in RD+I (open bars) versus HFD+I (filled bars) graft recipients at 3 weeks (D, E), 7 weeks (F, G), and 11 weeks (H, I) on diet. Each dot in D-I represents a single mouse. Data are mean ± SEM with individual points shown. *P < 0.05, **P < 0.01, ***P < 0.001.

**Figure S3. Islet transcription factor analysis of human grafts.** (A, B) Representative immunofluorescence for INS (green), MAFA (red), and DAPI (blue) in RD+I (black bars) and HFD+I (red bars) grafts, with quantification of the percentage of MAFA-positive beta-cells. (C, D) Representative immunofluorescence for GCG (green), ARX (red), and DAPI (blue), with quantification of the percentage of ARX-positive alpha-cells. Scale bar = 50 µm for all images in A and C. (E-I) CODEX quantification of the percentage of alpha- and beta-cells positive for PAX6 (E), PDX1 (F), ISL1 (G), IRX2 (H), and NEUROD1 (I) between RD+I and HFD+I grafts. Data are mean ± SEM with individual points shown.

**Figure S4. Graft amyloid and immune infiltration.** (A, B) Thioflavin S (green), INS (red), and DAPI (blue) staining showing increased amyloid in HFD+I (red bars) grafts. (C, D) Chromogranin A (CGA, green), CD45 (red), and DAPI (blue) in RD+I (black bars) and HFD+I (red bars) grafts. Scale bar = 200 µm for all images in A and C. Data are mean ± SEM with individual points shown. **P < 0.01.

**Figure S5. CODEX quantification of human-derived immune, vascular, and extracellular matrix markers in human islet grafts.** Percentage of marker-positive area relative to total graft area in RD+I (black bars) versus HFD+I (red bars) grafts for the immune and antigen-presenting markers CD68 (A), CD163 (B), MMR (C), and HLA-DR (D); the pericyte marker MCAM (E); the endothelial markers CD31 (F) and SELP (G); the extracellular matrix components laminin (H), collagen I (I), collagen IV (J), and vimentin (K); the epithelial marker keratin (L); the macrophage marker IBA1 (M); and collagen VI (N). Antibodies in this panel, including CD45 and CD31, are human-specific and therefore report on human-derived cells within the graft rather than on host mouse cells. No marker differed significantly between diet groups; collagen I showed a non-significant trend toward higher values in HFD+I grafts (I; P = 0.0565). Each dot represents a single graft. Data are mean ± SEM with individual points shown.

**Figure S6. Validation of the GKO-NSG-DTR ablation model and its sex-dependent response.** (A) Blood glucose in male GKO-NSG-DTR mice after DT injection (red arrow), showing consistent sustained hyperglycemia by day 3. (B) Blood glucose in DT non-responder female GKO-NSG-DTR mice. (C) Blood glucose in DT-responder female GKO-NSG-DTR mice, illustrating heterogeneous and often delayed hyperglycemia. (D) Representative immunofluorescence images of host pancreatic islets from female GKO-NSG-DTR mice after PBS or DT injection and (E) from male mice, including NSG, GKO-NSG-DTR, and human islet-transplanted GKO-NSG-DTR (PBS and DT) recipients, labeled for INS (green), Pro-GCG (red), and SST (blue). Scale bar = 50 µm for all images in D and E. Data are mean ± SEM with individual points shown.

**Figure S7. Human islet graft integrity after DT-mediated host beta-cell ablation.** (A) Representative gross images of graft-bearing kidneys at retrieval from PBS- and DT-treated GKO-NSG-DTR recipients (two independent pairs; ruler in cm). Arrows indicate the graft. (B) Representative whole-graft sections from PBS- and DT-treated recipients labeled for insulin (INS, green), proglucagon (Pro-GCG, red), and somatostatin (SST, blue). (C) Quantification of graft endocrine composition, expressed as the percentage of alpha-, beta-, and delta-cells per graft, in PBS- (black bars) and DT-treated (orange bars) recipients. Each dot represents a single graft. Data are mean ± SEM with individual points shown. *P < 0.05, **P < 0.01.

## Acknowledgments and Funding

We gratefully acknowledge the organ donors and their families for their invaluable gift, which made this research possible. We would also like to thank Greg Poffenberger for assistance with DT-related studies. Human pancreatic islets and/or other resources were provided by the NIDDK-funded Integrated Islet Distribution Program (IIDP) (RRID:SCR _014387) at City of Hope, NIH Grant # U24DK098085, the Human Pancreas Analysis Program^49–52^ (HPAP; RRID:SCR_016202; https://hpap.pmacs.upenn.edu/). HPAP is part of a Human Islet Research Network (HIRN; RRID:SCR_014393; https://hirnetwork.org/) with funding from DK112217, DK123594, DK112232, and DK123716. This work was supported by the U.S. Department of Veterans Affairs (IK2BX006210 [KCC] and BX000666 [ACP]), the National Institute of Diabetes and Digestive and Kidney Diseases (DK112217, DK123594, DK106755, DK112232, DK123716 [ACP] and 1 UG3 DK142192 [LDS]), the Vanderbilt Diabetes Research and Training Center (DK020593), Breakthrough T1D (4-COE-2025-1751-A-N [LDS]), and The Leona M. and Harry B. Helmsley Charitable Trust.

## Data Availability Statement

The datasets generated and analyzed during the current study are available from the corresponding authors on reasonable request.

## References

1. Faerch K, Vaag A, Holst JJ, Glumer C, Pedersen O, Borch-Johnsen K. Impaired fasting glycaemia vs impaired glucose tolerance: similar impairment of pancreatic alpha and beta cell function but differential roles of incretin hormones and insulin action. Diabetologia 2008;51(5):853–61. DOI: 10.1007/s00125-008-0951-x.

2. Newgard CB, An J, Bain JR, et al. A branched-chain amino acid-related metabolic signature that differentiates obese and lean humans and contributes to insulin resistance. Cell Metab 2009;9(4):311–26. DOI: 10.1016/j.cmet.2009.02.002.

3. Starke AAR, Erhardt G, Berger M, Zimmermann H. Elevated Pancreatic Glucagon in Obesity. Diabetes 1984;33(3):277–280. DOI: 10.2337/diab.33.3.277.

4. Krssak M, Brehm A, Bernroider E, et al. Alterations in postprandial hepatic glycogen metabolism in type 2 diabetes. Diabetes 2004;53(12):3048–56. DOI: 10.2337/diabetes.53.12.3048.

5. Hwang JH, Perseghin G, Rothman DL, et al. Impaired net hepatic glycogen synthesis in insulin-dependent diabetic subjects during mixed meal ingestion. A 13C nuclear magnetic resonance spectroscopy study. J Clin Invest 1995;95(2):783–7. DOI: 10.1172/JCI117727.

6. D’Alessio D. The role of dysregulated glucagon secretion in type 2 diabetes. Diabetes Obes Metab 2011;13 Suppl 1:126–32. DOI: 10.1111/j.1463-1326.2011.01449.x.

7. Unger RH. Glucagon Physiology and Pathophysiology. The New England journal of medicine 1971;285(8):443–449. DOI: 10.1056/NEJM197108192850806.

8. Baron AD, Schaeffer L, Shragg P, Kolterman OG. Role of hyperglucagonemia in maintenance of increased rates of hepatic glucose output in type II diabetics. Diabetes 1987;36(3):274–83. DOI: 10.2337/diab.36.3.274.

9. Reaven GM, Chen YD, Golay A, Swislocki AL, Jaspan JB. Documentation of hyperglucagonemia throughout the day in nonobese and obese patients with noninsulin-dependent diabetes mellitus. J Clin Endocrinol Metab 1987;64(1):106–10. DOI: 10.1210/jcem-64-1-106.

10. Unger RH, Aguilar-Parada E, Müller WA, Eisentraut AM. Studies of pancreatic alpha cell function in normal and diabetic subjects. The Journal of Clinical Investigation 1970;49(4):837–848. DOI: 10.1172/JCI106297.

11. Muller WA, Faloona GR, Aguilar-Parada E, Unger RH. Abnormal alpha-cell function in diabetes. Response to carbohydrate and protein ingestion. N Engl J Med 1970;283(3):109–15. DOI: 10.1056/NEJM197007162830301.

12. Unger R, Orci L. THE ESSENTIAL ROLE OF GLUCAGON IN THE PATHOGENESIS OF DIABETES MELLITUS. The Lancet 1975;305(7897):14–16. DOI: 10.1016/S0140-6736(75)92375-2.

13. Unger RH, Cherrington AD. Glucagonocentric restructuring of diabetes: a pathophysiologic and therapeutic makeover. J Clin Invest 2012;122(1):4–12. DOI: 10.1172/JCI60016.

14. Bolli G, de Feo P, Compagnucci P, et al. Abnormal glucose counterregulation in insulin-dependent diabetes mellitus. Interaction of anti-insulin antibodies and impaired glucagon and epinephrine secretion. Diabetes 1983;32(2):134–41. DOI: 10.2337/diab.32.2.134.

15. Gerich JE, Langlois M, Noacco C, Karam JH, Forsham PH. Lack of glucagon response to hypoglycemia in diabetes: evidence for an intrinsic pancreatic alpha cell defect. Science 1973;182(4108):171–3. DOI: 10.1126/science.182.4108.171.

16. Brissova M, Haliyur R, Saunders D, et al. alpha Cell Function and Gene Expression Are Compromised in Type 1 Diabetes. Cell Rep 2018;22(10):2667–2676. DOI: 10.1016/j.celrep.2018.02.032.

17. Adams JD, Dalla Man C, Laurenti MC, et al. Fasting glucagon concentrations are associated with longitudinal decline of β-cell function in non-diabetic humans. Metabolism 2020;105:154175. (In eng). DOI: 10.1016/j.metabol.2020.154175.

18. Adams JD, Egan AM, Laurenti MC, et al. Insulin secretion and action and the response of endogenous glucose production to a lack of glucagon suppression in nondiabetic subjects. Am J Physiol Endocrinol Metab 2021;321(5):E728–e736. (In eng). DOI: 10.1152/ajpendo.00284.2021.

19. Chang L, Kong L, Yan S, et al. Alpha cell dysfunction in type 2 diabetes: associations with insulin resistance and reduced insulin secretion. BMJ Open Diabetes Res Care 2026;14(3) (In eng). DOI: 10.1136/bmjdrc-2025-005653.

20. Chang L, Liu Y, Gu Y, et al. Inadequate Glucagon Suppression During OGTT in Prediabetes: A Systematic Review and Meta-analysis. J Clin Endocrinol Metab 2024;109(10):2673–2680. (In eng). DOI: 10.1210/clinem/dgae460.

21. Kohlenberg JD, Laurenti MC, Egan AM, et al. Differential contribution of alpha and beta cell dysfunction to impaired fasting glucose and impaired glucose tolerance. Diabetologia 2023;66(1):201–212. (In eng). DOI: 10.1007/s00125-022-05794-3.

22. Mohan S, Christie HE, Laurenti MC, et al. Abnormal Glucagon Secretion Contributes to a Longitudinal Decline in Glucose Tolerance. J Clin Endocrinol Metab 2025;110(9):e2956–e2965. (In eng). DOI: 10.1210/clinem/dgae915.

23. Salehi M. Alpha-Cell Secretion Across the Spectrum of Glucose Tolerance. J Clin Endocrinol Metab 2024;109(6):e1456–e1457. (In eng). DOI: 10.1210/clinem/dgad686.

24. El K, Capozzi ME, Campbell JE. Repositioning the Alpha Cell in Postprandial Metabolism. Endocrinology 2020;161(11) (In eng). DOI: 10.1210/endocr/bqaa169.

25. Deng B, Ruan T, Lu W, et al. Safety and efficacy of GLP-1 and glucagon receptor dual agonist for the treatment of type 2 diabetes and obesity: a systematic review and meta-analysis of randomized controlled trials. Endocrine 2024;86(1):15–27. (In eng). DOI: 10.1007/s12020-024-03857-6.

26. Capozzi ME, D’Alessio DA, Campbell JE. The past, present, and future physiology and pharmacology of glucagon. Cell Metab 2022;34(11):1654–1674. (In eng). DOI: 10.1016/j.cmet.2022.10.001.

27. Novikoff A, Müller TD. The molecular pharmacology of glucagon agonists in diabetes and obesity. Peptides 2023;165:171003. (In eng). DOI: 10.1016/j.peptides.2023.171003.

28. Pettus J, Boeder SC, Christiansen MP, et al. Glucagon receptor antagonist volagidemab in type 1 diabetes: a 12-week, randomized, double-blind, phase 2 trial. Nat Med 2022;28(10):2092–2099. (In eng). DOI: 10.1038/s41591-022-02011-x.

29. Rodriguez-Diaz R, Abdulreda MH, Formoso AL, et al. Innervation patterns of autonomic axons in the human endocrine pancreas. Cell Metab 2011;14(1):45–54. DOI: 10.1016/j.cmet.2011.05.008.

30. Richardson TM, Saunders DC, Haliyur R, et al. Human pancreatic capillaries and nerve fibers persist in type 1 diabetes despite beta cell loss. Am J Physiol Endocrinol Metab 2023;324(3):E251–E267. DOI: 10.1152/ajpendo.00246.2022.

31. Brissova M, Fowler MJ, Nicholson WE, et al. Assessment of human pancreatic islet architecture and composition by laser scanning confocal microscopy. J Histochem Cytochem 2005;53(9):1087–97. DOI: 10.1369/jhc.5C6684.2005.

32. Dai C, Brissova M, Hang Y, et al. Islet-enriched gene expression and glucose-induced insulin secretion in human and mouse islets. Diabetologia 2012;55(3):707–18. DOI: 10.1007/s00125-011-2369-0.

33. Cabrera O, Berman DM, Kenyon NS, Ricordi C, Berggren PO, Caicedo A. The unique cytoarchitecture of human pancreatic islets has implications for islet cell function. Proc Natl Acad Sci U S A 2006;103(7):2334–9. DOI: 10.1073/pnas.0510790103.

34. Dai C, Kayton NS, Shostak A, et al. Stress-impaired transcription factor expression and insulin secretion in transplanted human islets. J Clin Invest 2016;126(5):1857–70. DOI: 10.1172/JCI83657.

35. Brozzi F, Nardelli TR, Lopes M, et al. Cytokines induce endoplasmic reticulum stress in human, rat and mouse beta cells via different mechanisms. Diabetologia 2015;58(10):2307–16. DOI: 10.1007/s00125-015-3669-6.

36. Arda HE, Li L, Tsai J, et al. Age-Dependent Pancreatic Gene Regulation Reveals Mechanisms Governing Human beta Cell Function. Cell Metab 2016;23(5):909–20. DOI: 10.1016/j.cmet.2016.04.002.

37. Conrad E, Dai C, Spaeth J, et al. The MAFB transcription factor impacts islet alpha-cell function in rodents and represents a unique signature of primate islet beta-cells. Am J Physiol Endocrinol Metab 2016;310(1):E91–E102. DOI: 10.1152/ajpendo.00285.2015.

38. Conrad E, Stein R, Hunter CS. Revealing transcription factors during human pancreatic beta cell development. Trends Endocrinol Metab 2014;25(8):407–14. DOI: 10.1016/j.tem.2014.03.013.

39. Cyphert HA, Walker EM, Hang Y, et al. Examining How the MAFB Transcription Factor Affects Islet beta-Cell Function Postnatally. Diabetes 2019;68(2):337–348. DOI: 10.2337/db18-0903.

40. Walker JT, Saunders DC, Brissova M, Powers AC. The Human Islet: Mini-Organ With Mega-Impact. Endocr Rev 2021;42(5):605–657. DOI: 10.1210/endrev/bnab010.

41. Evans-Molina C, Pettway YD, Saunders DC, et al. Heterogeneous endocrine cell composition defines human islet functional phenotypes. Nat Commun 2026;17(1). DOI: 10.1038/s41467-026-70689-5.

42. Dai XQ, Camunas-Soler J, Briant LJB, et al. Heterogenous impairment of alpha cell function in type 2 diabetes is linked to cell maturation state. Cell Metab 2022;34(2):256–268 e5. DOI: 10.1016/j.cmet.2021.12.021.

43. Zhang Q, Ramracheya R, Lahmann C, et al. Role of KATP Channels in Glucose-Regulated Glucagon Secretion and Impaired Counterregulation in Type 2 Diabetes. Cell Metabolism 2013;18(6):871–882. DOI: 10.1016/j.cmet.2013.10.014.

44. Omar-Hmeadi M, Lund PE, Gandasi NR, Tengholm A, Barg S. Paracrine control of alpha-cell glucagon exocytosis is compromised in human type-2 diabetes. Nat Commun 2020;11(1):1896. DOI: 10.1038/s41467-020-15717-8.

45. Walker JT, Saunders DC, Rai V, et al. Genetic risk converges on regulatory networks mediating early type 2 diabetes. Nature 2023;624(7992):621–629. DOI: 10.1038/s41586-023-06693-2.

46. Surwit RS, Kuhn CM, Cochrane C, McCubbin JA, Feinglos MN. Diet-Induced Type II Diabetes in C57BL/6J Mice. Diabetes 1988;37(9):1163–1167. DOI: 10.2337/diab.37.9.1163.

47. Kellard JA, Rorsman NJG, Hill TG, et al. Reduced somatostatin signalling leads to hypersecretion of glucagon in mice fed a high-fat diet. Molecular Metabolism 2020;40:101021. DOI: 10.1016/j.molmet.2020.101021.

48. Tellez K, Hang Y, Gu X, Chang CA, Stein RW, Kim SK. In vivo studies of glucagon secretion by human islets transplanted in mice. Nat Metab 2020;2(6):547–557. DOI: 10.1038/s42255-020-0213-x.

49. Kaestner KH, Powers AC, Naji A, Consortium H, Atkinson MA. NIH Initiative to Improve Understanding of the Pancreas, Islet, and Autoimmunity in Type 1 Diabetes: The Human Pancreas Analysis Program (HPAP). Diabetes 2019;68(7):1394–1402. DOI: 10.2337/db19-0058.

50. Shapira SN, Naji A, Atkinson MA, Powers AC, Kaestner KH. Understanding islet dysfunction in type 2 diabetes through multidimensional pancreatic phenotyping: The Human Pancreas Analysis Program. Cell Metab 2022;34(12):1906–1913. DOI: 10.1016/j.cmet.2022.09.013.

51. Fasolino M, Schwartz GW, Patil AR, et al. Single-cell multi-omics analysis of human pancreatic islets reveals novel cellular states in type 1 diabetes. Nat Metab 2022;4(2):284–299. DOI: 10.1038/s42255-022-00531-x.

52. Patil AR, Schug J, Naji A, Kaestner KH, Faryabi RB, Vahedi G. Single-cell expression profiling of islets generated by the Human Pancreas Analysis Program. Nat Metab 2023;5(5):713–715. DOI: 10.1038/s42255-023-00806-x.

53. Hart NJ, Powers AC. Use of human islets to understand islet biology and diabetes: progress, challenges and suggestions. Diabetologia 2019;62(2):212–222. DOI: 10.1007/s00125-018-4772-2.

54. Poitout V, Satin LS, Kahn SE, et al. A call for improved reporting of human islet characteristics in research articles. Diabetologia 2019;62(2):209–211. (In eng). DOI: 10.1007/s00125-018-4784-y.

55. Coate KC, Dai C, Singh A, et al. Interruption of glucagon signaling augments islet non-alpha cell proliferation in SLC7A2- and mTOR-dependent manners. Mol Metab 2024;90:102050. DOI: 10.1016/j.molmet.2024.102050.

56. Dean ED, Li M, Prasad N, et al. Interrupted Glucagon Signaling Reveals Hepatic alpha Cell Axis and Role for L-Glutamine in alpha Cell Proliferation. Cell Metab 2017;25(6):1362–1373 e5. DOI: 10.1016/j.cmet.2017.05.011.

57. Marzban L, Park K, Verchere CB. Islet amyloid polypeptide and type 2 diabetes. Exp Gerontol 2003;38(4):347–51. (In eng). DOI: 10.1016/s0531-5565(03)00004-4.

58. Westermark P, Andersson A, Westermark GT. Islet amyloid polypeptide, islet amyloid, and diabetes mellitus. Physiol Rev 2011;91(3):795–826. (In eng). DOI: 10.1152/physrev.00042.2009.

59. Black S, Phillips D, Hickey JW, et al. CODEX multiplexed tissue imaging with DNA-conjugated antibodies. Nat Protoc 2021;16(8):3802–3835. DOI: 10.1038/s41596-021-00556-8.

60. Dai C, Zhang Y, Gong Y, et al. Hyperaminoacidemia from interrupted glucagon signaling increases pancreatic acinar cell proliferation and size via mTORC1 and YAP pathways. iScience 2024;27(12):111447. (In eng). DOI: 10.1016/j.isci.2024.111447.

61. Coate K, Liu J, Guo M, et al. MAFB is essential for the maintenance of adult human α-cell identity and glucagon secretion. bioRxiv 2026:2026.08.08.743687. DOI: 10.64898/2026.08.08.743687.

62. Aigha, II, Abdelalim EM. NKX6.1 transcription factor: a crucial regulator of pancreatic β cell development, identity, and proliferation. Stem Cell Res Ther 2020;11(1):459. (In eng). DOI: 10.1186/s13287-020-01977-0.

63. Brissova M, Shostak A, Fligner CL, et al. Human Islets Have Fewer Blood Vessels than Mouse Islets and the Density of Islet Vascular Structures Is Increased in Type 2 Diabetes. J Histochem Cytochem 2015;63(8):637–45. DOI: 10.1369/0022155415573324.

64. Meier DT, Morcos M, Samarasekera T, Zraika S, Hull RL, Kahn SE. Islet amyloid formation is an important determinant for inducing islet inflammation in high-fat-fed human IAPP transgenic mice. Diabetologia 2014;57(9):1884–8. DOI: 10.1007/s00125-014-3304-y.

65. Shrestha S, Jennings LT, Knofczynski K, Shivakumar SB, Peterson QP. Modeling diabetic alpha cell dysfunction using stem cell-derived alpha cells. Stem Cell Reports 2025;20(6):102504. DOI: 10.1016/j.stemcr.2025.102504.

66. Guo S, Dai C, Guo M, et al. Inactivation of specific β cell transcription factors in type 2 diabetes. J Clin Invest 2013;123(8):3305–16. (In eng). DOI: 10.1172/jci65390.

67. Dobbs R, Sakurai H, Sasaki H, et al. Glucagon: Role in the Hyperglycemia of Diabetes Mellitus. Science 1975;187(4176):544–547. DOI: doi:10.1126/science.1089999.

68. Artner I, Hang Y, Mazur M, et al. MafA and MafB regulate genes critical to beta-cells in a unique temporal manner. Diabetes 2010;59(10):2530–9. DOI: 10.2337/db10-0190.

69. Cataldo LR, Singh T, Achanta K, et al. MAFA and MAFB regulate exocytosis-related genes in human beta-cells. Acta Physiol (Oxf) 2022;234(2):e13761. DOI: 10.1111/apha.13761.

70. Gao R, Acreman S, Ma J, Abdulkader F, Wendt A, Zhang Q. α-cell electrophysiology and the regulation of glucagon secretion. J Endocrinol 2023;258(2) (In eng). DOI: 10.1530/joe-22-0295.

71. Bosi E, Marchetti P, Rutter GA, Eizirik DL. Human alpha cell transcriptomic signatures of types 1 and 2 diabetes highlight disease-specific dysfunction pathways. iScience 2022;25(10):105056. DOI: 10.1016/j.isci.2022.105056.

72. Rodriguez-Diaz R, Molano RD, Weitz JR, et al. Paracrine Interactions within the Pancreatic Islet Determine the Glycemic Set Point. Cell Metab 2018;27(3):549–558.e4. (In eng). DOI: 10.1016/j.cmet.2018.01.015.

73. Wang YJ, Traum D, Schug J, et al. Multiplexed In Situ Imaging Mass Cytometry Analysis of the Human Endocrine Pancreas and Immune System in Type 1 Diabetes. Cell Metab 2019;29(3):769–783.e4. (In eng). DOI: 10.1016/j.cmet.2019.01.003.

74. Wu M, Lee MYY, Bahl V, et al. Single-cell analysis of the human pancreas in type 2 diabetes using multi-spectral imaging mass cytometry. Cell Rep 2021;37(5):109919. (In eng). DOI: 10.1016/j.celrep.2021.109919.

