## Supplementary material for "High-fat feeding increases graft-derived glucagon output and impairs beta-cell secretion in transplanted human islets": Supp Figures

### Fig. S1

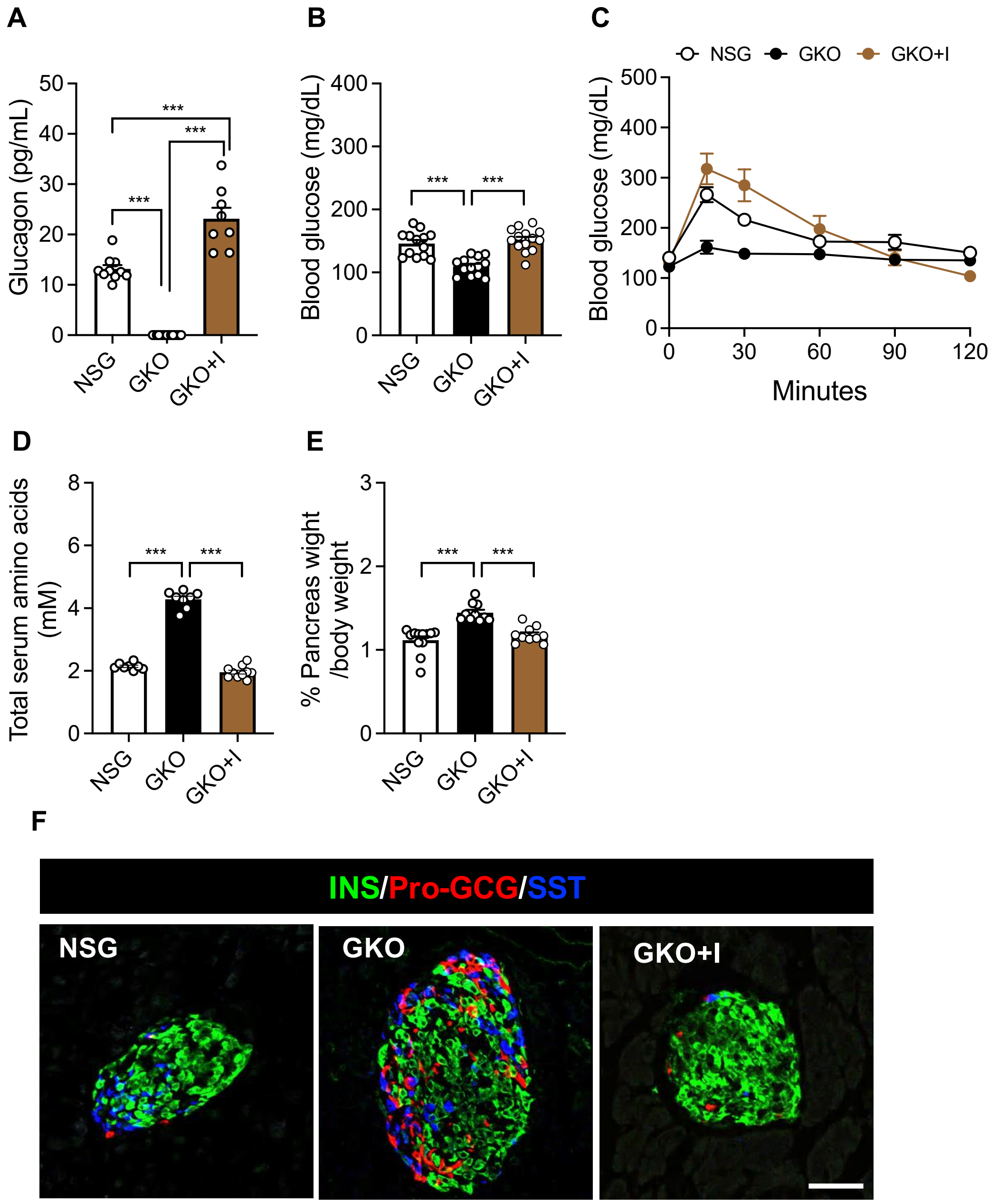
A
Glucagon (pg/mL)
NSG
GKO
GKO+I
B
Blood glucose (mg/dL)
C
NSG GKO GKO+I
Minutes
D
Total serum amino acids (mM)
E
% Pancreas wight /body weight
F
INS/Pro-GCG/SST
NSG
GKO
GKO+I

**Fig. S2**

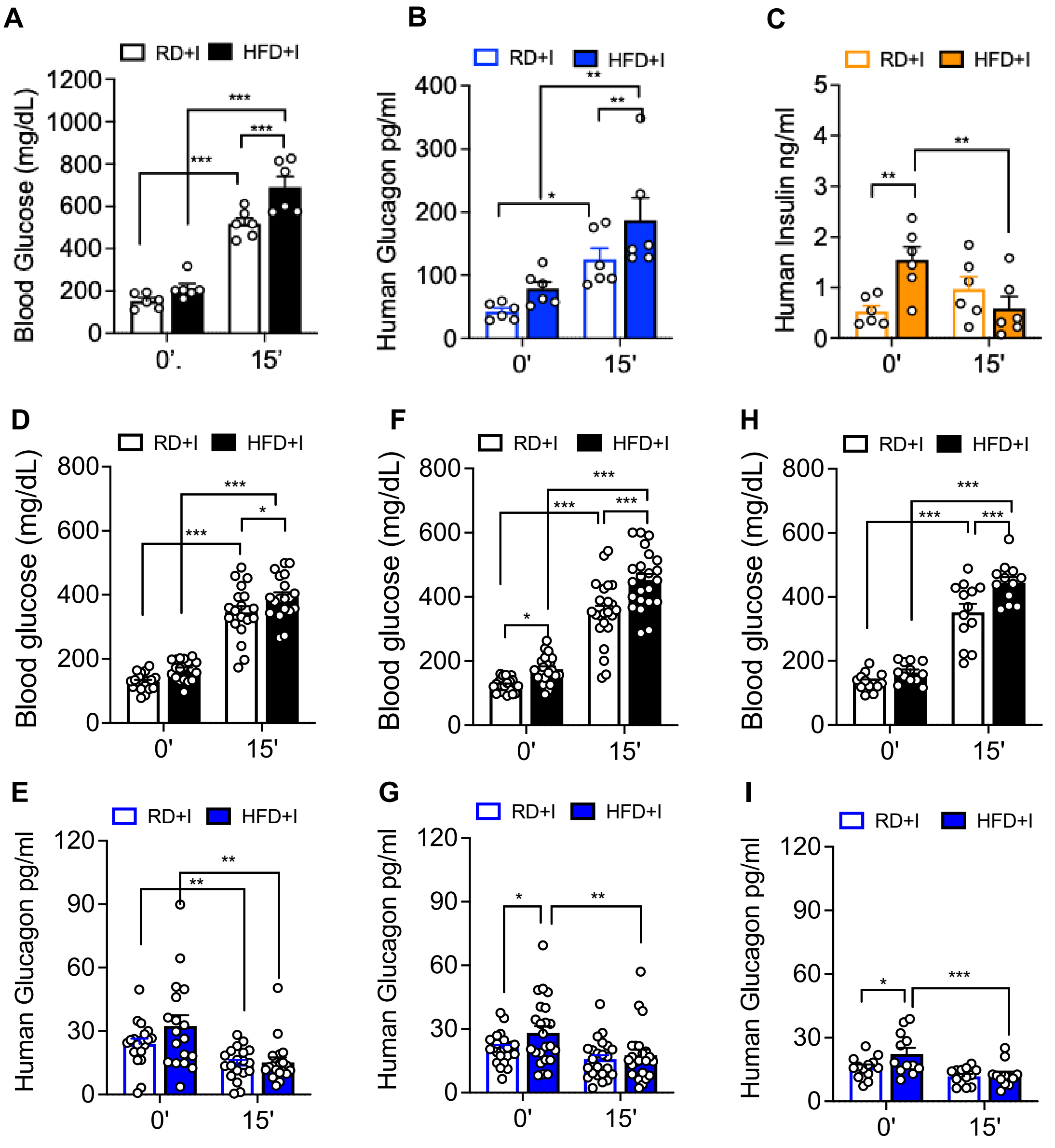

A
RD+I HFD+I
Blood Glucose (mg/dL)
0'. 15'
B
RD+I HFD+I
Human Glucagon pg/ml
0' 15'
C
RD+I HFD+I
Human Insulin ng/ml
0' 15'
D
RD+I HFD+I
Blood glucose (mg/dL)
0' 15'
F
RD+I HFD+I
Blood glucose (mg/dL)
0' 15'
H
RD+I HFD+I
Blood glucose (mg/dL)
0' 15'
E
RD+I HFD+I
Human Glucagon pg/ml
0' 15'
G
RD+I HFD+I
Human Glucagon pg/ml
0' 15'
I
RD+I HFD+I
Human Glucagon pg/ml
0' 15'

Fig. S3

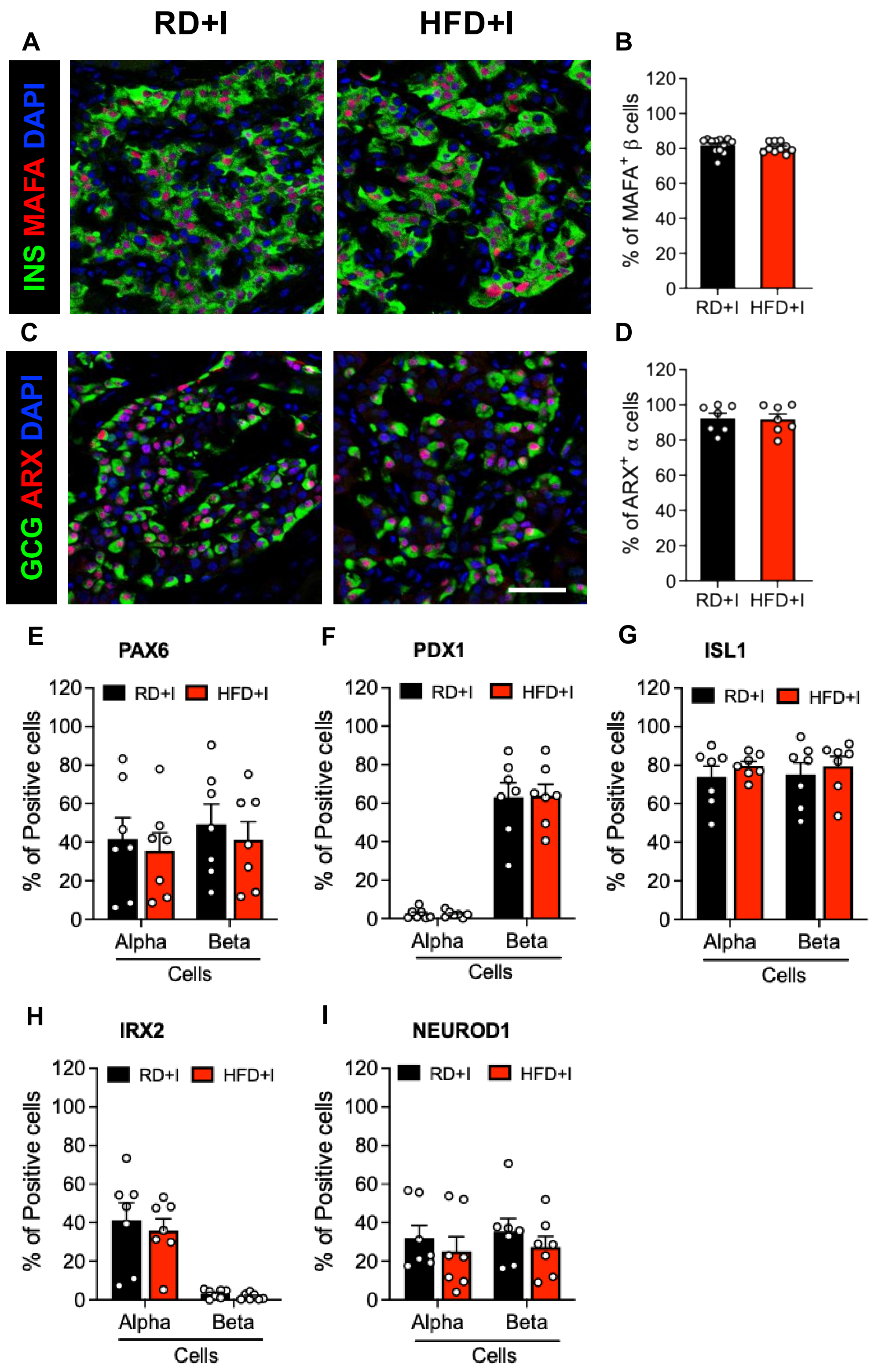

A
RD+I
HFD+I
INS MAFA DAPI
B
% of MAFA+ β cells
RD+I HFD+I
C
GCG ARX DAPI
D
% of ARX+ α cells
RD+I HFD+I
E
PAX6
F
PDX1
G
ISL1
H
IRX2
I
NEUROD1
RD+I HFD+I
% of Positive cells
Alpha Beta
Cells

**Fig. S4**

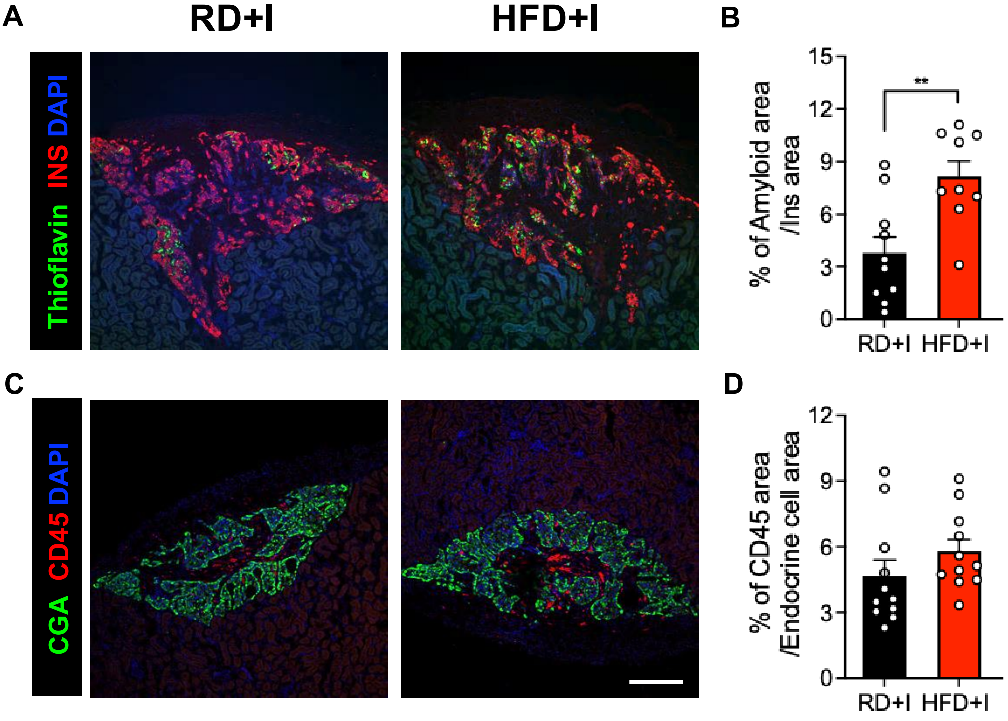
A
RD+I
HFD+I
Thioflavin INS DAPI
B
% of Amyloid area /Ins area
15
12
9
6
3
0
**
RD+I
HFD+I
C
CGA CD45 DAPI
D
% of CD45 area /Endocrine cell area
12
9
6
3
0
RD+I
HFD+I

**Fig. S5**

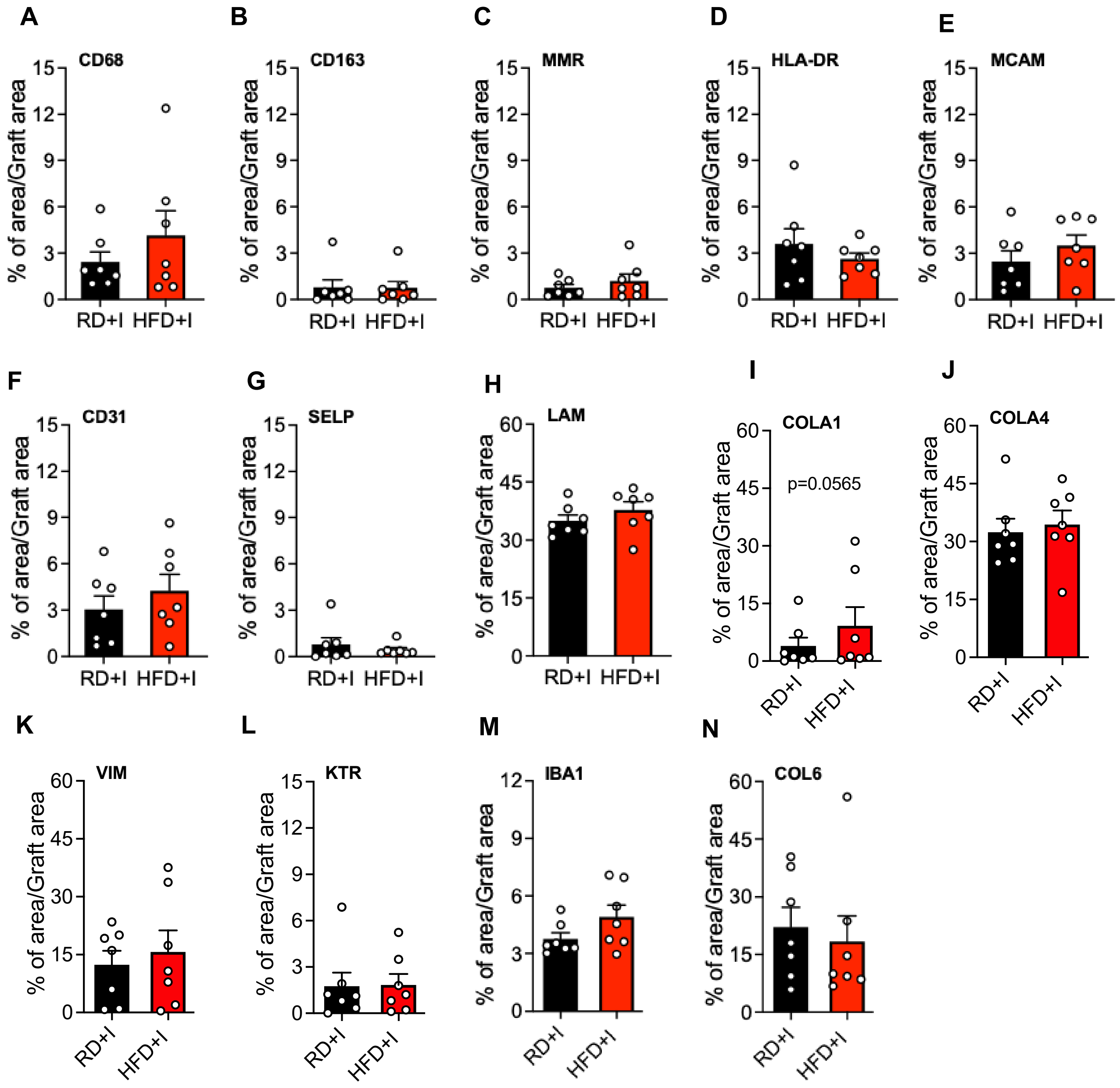
A
CD68
B
CD163
C
MMR
D
HLA-DR
E
MCAM
F
CD31
G
SELP
H
LAM
I
COLA1
p=0.0565
J
COLA4
K
VIM
L
KTR
M
IBA1
N
COL6
% of area/Graft area
RD+I
HFD+I

Fig. S6

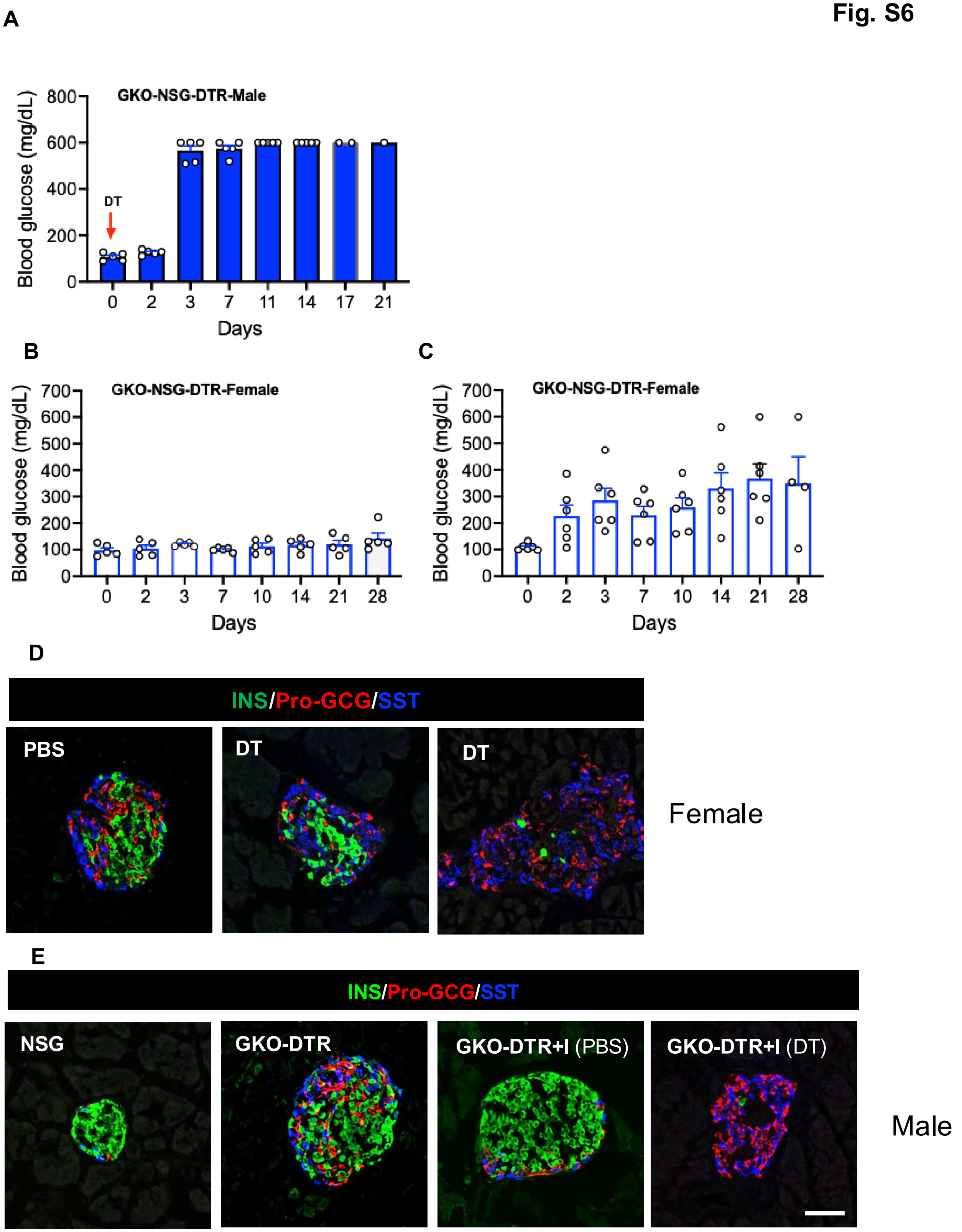

A
GKO-NSG-DTR-Male
Blood glucose (mg/dL)
800
600
400
200
0
DT
0
2
3
7
11
14
17
21
Days
B
GKO-NSG-DTR-Female
Blood glucose (mg/dL)
700
600
500
400
300
200
100
0
0
2
3
7
10
14
21
28
Days
C
GKO-NSG-DTR-Female
Blood glucose (mg/dL)
700
600
500
400
300
200
100
0
0
2
3
7
10
14
21
28
Days
D
INS/Pro-GCG/SST
PBS
DT
DT
Female
E
INS/Pro-GCG/SST
NSG
GKO-DTR
GKO-DTR+I (PBS)
GKO-DTR+I (DT)
Male

**Fig. S7**

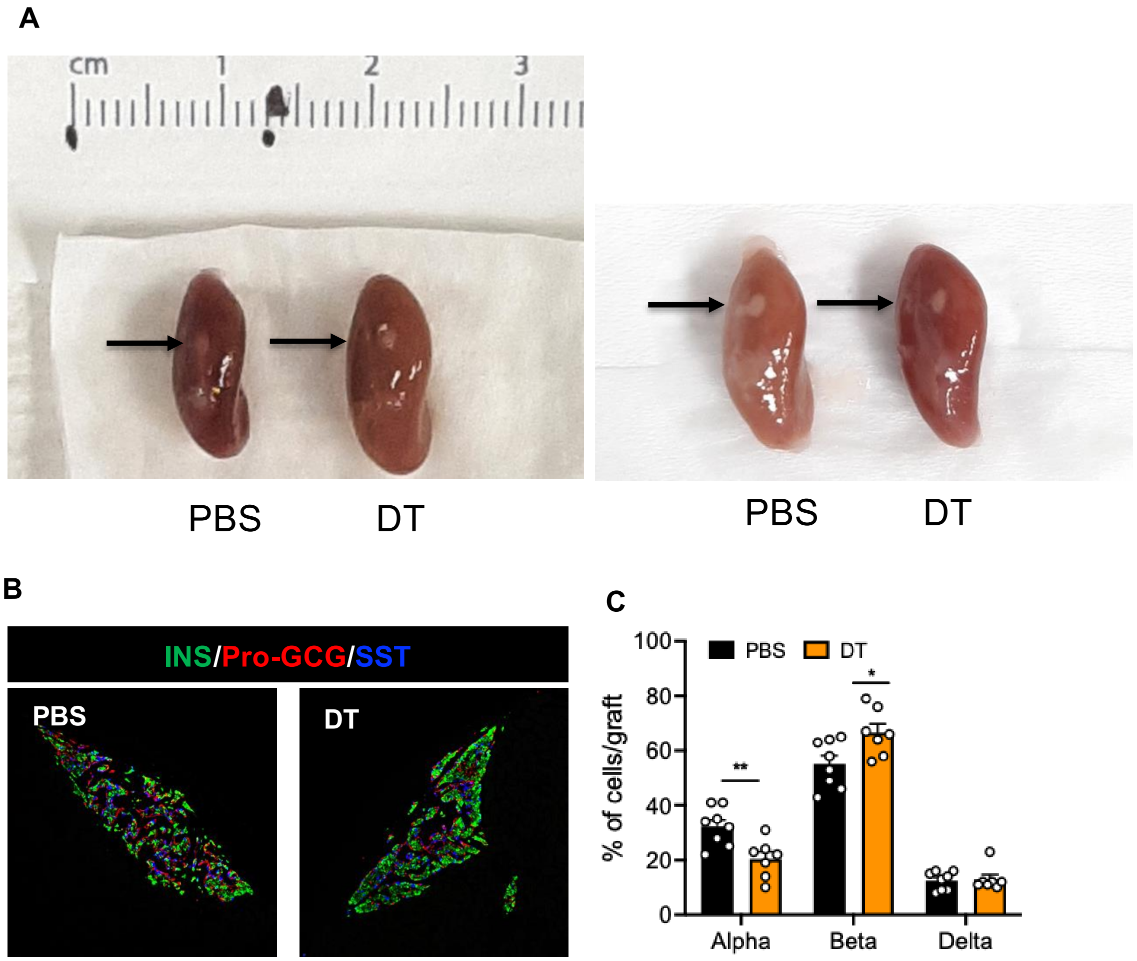
A
cm
1
2
3
PBS
DT
PBS
DT
B
INS/Pro-GCG/SST
PBS
DT
C
PBS
DT
% of cells/graft
100
80
60
40
20
0
**
*
Alpha
Beta
Delta
