## Supplementary material for "High-fat feeding increases graft-derived glucagon output and impairs beta-cell secretion in transplanted human islets": Supp Table 1

**Supplemental Table 1: Human islet preparations used in this study**

| Donor | Unique ID | Age years | Sex | BMI (kg/m2) | HbA1C (%) | Ethnicity/Race | Cause of Death |
|---|---|---|---|---|---|---|---|
| 1 | SAMN28088753 | 41 | F | 31.6 | 5.3% | Hispanic | Anoxia |
| 2 | AJFL134 * | 55 | F | 28.6 | 5.9 | N/A | COD |
| 3 | SAMN31525423 | 30 | F | 39.90 | N/A | Black | N/A |
| 4 | SAMN31896841 | 66 | F | 27.4 | 5.4% | N/A | N/A |
| 5 | AKCB359 | 51 | F | 43.7 | 5.6% | Black | Anoxia |
| 6 | SAMN35133168 | 50 | F | 29.70 | 5.4% | Hispanic | COD |
| 7 | AIHQ341 | 4 | M | 20.63 | 4.9% | Hispanic | Anoxia |
| | AJFL134 * | 55 | F | 28.6 | 5.9 | N/A | COD |
| 8 | AJHI084 | 19 | F | 18.1 | 5.2% | Hispanic | N/A |
| 9 | AJKM384 | 29 | M | 27.24 | 5.4% | Caucasian | N/A |

Abbreviations: HPAP, Human Pancreas Analysis Program; IIAM, International Institute for the A
IDP, Integrated Islet Distribution Program; IP, Imagine Pharma; N/A, Not Availabl

* Same donor for two separate transplantations

All donors did not have history of diabetes.

All islets were handpicked to purity >95%

| Islet Source | Experiment | # of mice transplanted RD/HFD or PBS/DT |
|---|---|---|
| HPAP | HFD | 3, 3 |
| IIAM | HFD | 4, 4 |
| IIDP | HFD | 5, 5 |
| IIDP | HFD | 4, 4 |
| IIAM | HFD | 4, 4 |
| IIDP | HFD | 3, 3 |
| HPAP | DTR | 4, 4 |
| IIAM | DTR | 2, 2 |
| HPAP | DTR | 3, 3 |
| HPAP | DTR | 3, 3 |

dvancement of Medicine;
le
