## Supplementary material for "High-fat feeding increases graft-derived glucagon output and impairs beta-cell secretion in transplanted human islets": Supp Table 2

**Supplemental Table 2: Primary antibodies and secondary antibodies**

| Antigen | Host | Class | Source | Catalog # |
|---|---|---|---|---|
| ARX | Sheep | Polyclonal | R&D | AF7068 |
| CD45 | Mouse | Monoclonal | Fisher scientific/BD | BD347460 |
| Chromogranin A | Rabit | Polyclonal | R&D / Novus | NB-120-15160 |
| cPeptide | Rat | Monoclonal | DSHB | GN-ID4 |
| Glucagon | Guinea pig | Polyclonal | LSBio | LS-C202759 |
| Glucagon | Rabit | polyclonal | Cell Signaling | 2760 |
| Insulin | Guinea pig | polyclonal | Dako | A05564 |
| MAFA | Rabit | Monoclonal | Bethyl | A700-067 |
| MAFB | Mouse | Monoclonal | R&D | MAB 3810 |
| NKX6.1 | Mouse | Monoclonal | BD Pharmingen | 563022 |
| Pro-glucagon | Rabit | Monoclonal | Cell Signaling | 8233 |
| Somatostatin | Goat | Polyclonal | Santa Cruz | sc-7819 |
| **Secondary Antibodies** | | | | |
| Rabbit | Donkey | cy2 | Jackson Labs | 711-225-152 |
| Rabbit | Donkey | cy3 | Jackson Labs | 711-165-152 |
| Rabbit | Donkey | cy5 | Jackson Labs | 711-175-152 |
| Mouse | Donkey | cy3 | Jackson Labs | 315-165-045 |
| Goat | Donkey | cy5 | Jackson Labs | 705-175-147 |
| Rat | Donkey | cy2 | Jackson Labs | 712-225-153 |
| Sheep | Donkey | cy3 | Jackson Labs | 713-165-147 |
| Guinea Pig | Donkey | cy2 | Jackson Labs | 706-225-148 |
| Guinea Pig | Donkey | cy3 | Jackson Labs | 706-165-148 |
| Guinea Pig | Donkey | cy5 | Jackson Labs | 706-175-148 |
