## Supplementary material for "High-fat feeding increases graft-derived glucagon output and impairs beta-cell secretion in transplanted human islets": Supp Table 3

**Supplemental Table 3: Antibodies used in CODEX**

| Marker name | RX | Type | Clone |
|---|---|---|---|
| CD45 | 488 | Custom, Akoya | HI30 |
| NEUROD1 | 488 | Leinco | R8-294 |
| HLA-DR | 488 | Custom | L243 |
| COL1A1 | 488 | Custom | COL-1 |
| KRT | 488 | Akoya | AE-1/AE-3 |
| SELP | 488 | Leinco | Polyclonal |
| CPEP | 488 | Custom | C-PEP-01 |
| CD163 | 488 | Custom | GHI/61 |
| MCAM | 488 | Custom | P1H12 |
| proINS | 488 | Leinco | GS-9A8 |
| GCG | 550 | Custom | K79bB10 |
| IRX2 | 550 | Leinco | 1B7 |
| PDX1 | 550 | Leinco | D59H3 |
| SST | 550 | Custom | 7G5 |
| PAX6 | 550 | Leinco | Polyclonal |
| NKX6-1 | 550 | Custom | R11-560 |
| CD31 | 550 | Akoya, Custom | WM59 |
| SYP | 550 | Custom | 7H12 |
| COL6 | 550 | Custom | EPR17072 |
| ARX | 647 | Leinco | Polyclonal |
| LAM | 647 | Custom | Polyclonal |
| CHGA | 647 | Custom | LK2H10+PHE5+CGA/414 |
| PPY | 647 | Custom | 548416 |
| MMR | 647 | Custom | 15-2 |
| VIM | 647 | Custom | RV202 |
| COL4A1 | 647 | Custom | Polyclonal |
| IBA1 | 647 | Custom | EPR16589 |
| ISL1 | 647 | Leinco | Q11-465 |
| CD68 | 647 | Custom | KP1 |
